# Cationic charge and polyspecificity of an integrin domain regulates infectivity of malaria parasites

**DOI:** 10.1101/421800

**Authors:** Dennis Klug, Sarah Goellner, Julia Sattler, Leanne Strauss, Jessica Kehrer, Konrad Beyer, Miriam Reinig, Mirko Singer, Chafen Lu, Timothy A. Springer, Friedrich Frischknecht

## Abstract

Cell-cell and cell-substrate adhesion is critical for many functions in life. In eukaryotes, I-domains mediate functions as divergent as tissue traversal by malaria-causing *Plasmodium* parasites as well as cell adhesion and migration by human leucocytes. The I-domain containing protein TRAP is important for *Plasmodium* sporozoite motility and invasion. Here we show that the I-domain of TRAP is required to mediate adhesional properties which can be partially preserved when the native I-domain is replaced by I-domains from human integrins or from an apicomplexan parasite that does not infect insects. By putting in vivo data and structural features in perspective we conclude that polyspecificity and positive charge around the ligand binding site of the I-domain are important for TRAP function. Our data suggest a highly preserved functionality of I-domains across eukaryotic evolution that is used by apicomplexan parasites to invade a broad range of tissues in a variety of hosts.

## Introduction

Domains with similar overall structures, initially described for von Willebrand factor A-domains, are found in cell-surface proteins including integrins, extracellular matrix, and complement components, and mediate a diversity of functions including cell adhesion, migration, and signaling (Whittaker & Hynes 2002). Here, we study a subset of A-domains termed I-domains because they differ from von Willebrand factor A-domains in the position of their ligand binding site and in the presence of a metal ion-dependent adhesion site (MIDAS) within the ligand binding center (Liddington 2014). I-domains within integrins switch between open and closed states coordinately with conformational change in neighboring integrin domains. This switch from an open to a closed conformation in the I-domain around the MIDAS increases ligand-binding affinity by ∼1,000-fold (Schürpf & Springer 2011). Furthermore, I-domains are key modules in adhesins employed by apicomplexan pathogens. I-domain-containing, membrane spanning surface glycoproteins have been shown to be essential for tissue traversal and cell invasion by *Toxoplasma gondii* and *Plasmodium spp.* and are present in all known apicomplexans (Sultan et al. 1997; Morahan et al. 2009). In *Plasmodium*, a six I-domain containing protein named CTRP is required for invasion of the mosquito midgut by the ookinete (Mathias et al. 2013; Ramakrishnan et al. 2011). Once the ookinete crosses the midgut epithelium it forms an oocyst wherein it differentiates into hundreds of sporozoites (Frischknecht & Matuschewski 2017). Sporozoites use active motility to egress from the oocyst into the hemolymph (Klug & Frischknecht 2017), and subsequently enter the salivary glands from where they can be transmitted back to a vertebrate host. Once deposited in the skin during a blood meal by an infected mosquito, sporozoites migrate rapidly to find and enter blood vessels (Amino et al. 2006). Within the blood stream parasites are passively transported to the liver where they infect hepatocytes and develop into liver stages. The subsequent blood stages cause the typical symptoms of malaria by triggering a massive immune response, clogging capillaries and lysing red blood cells (Cowman et al. 2016).

Sporozoites express two adhesins with I-domains, TRAP (thrombospondin related anonymous protein) and TLP (TRAP-like protein). These proteins are stored in secretory vesicles called micronemes, at the apical end of the highly polarized sporozoite (Tomley & Soldati 2001). After fusion of micronemes with the plasma membrane, TRAP and TLP are present on the plasma membrane, where they form a bridge between extracellular ligands and the membrane-subtending actin-myosin motor that drives gliding motility and invasion (Heintzelman 2015; Frischknecht & Matuschewski 2017). Deletion of *tlp* causes only a mild phenotype in tissue traversal while deletion of *trap* yields sporozoites that cannot move productively, fail completely to enter into salivary glands and are unable to infect mice if isolated from mosquitoes and injected intravenously (Sultan et al. 1997; Moreira et al. 2008; Hellmann et al. 2013; Quadt et al. 2016). Mutations of amino acids within the MIDAS motif of the single I-domain in TRAP decrease the capacity of sporozoites to enter salivary glands and liver cells as well as to infect mice (Wengelnik et al. 1999; Matuschewski et al. 2002). However, these mutant sporozoites were still able to migrate. This suggests that the MIDAS is important for ligand binding but not for productive motility.

Crystal structures of the N-terminal portion of TRAP in *Plasmodium spp*. and a TRAP orthologue in *Toxoplasma gondii*, micronemal protein 2 (MIC2), revealed the I-domain in both open and closed conformations in association with a thrombospondin type-I repeat domain (Song et al. 2012; Song & Springer 2014) (**Figure 1**). The apicomplexan I-domains structurally resemble I-domains found in integrin α-subunits (αI-domains) much more than I-domains in integrin β-subunits (βI-domains). Between the closed and open states of both apicomplexan I-domains and integrin αI-domains, the Mg^2+^ ion at the MIDAS similarly moves ∼2 Å closer to one coordinating sidechain and away from another, and this movement is linked to essentially identical pistoning of the C-terminal, α7-helix toward the “bottom” of the domain (**Figure 1A compared to 1B and Figure 1D compared to 1E**). The distance pistoned is equivalent to two turns of an α-helix. Uniquely in the apicomplexan I-domains, a segment N-terminal to the I-domain is disulfide linked to the last helical turn of the α7-helix in its closed conformation (**Figure 1A-C**). As this segment pistons out of contact with the remainder of the I-domain in the open conformation, the last two turns of the α7-helix with its cysteine unwind, the N-terminal segment with its cysteine moves in a similar direction, and these segments reshape to form a β-ribbon (**Figure 1A**). Because of close structural homology between human integrin αI-and apicomplexan adhesin I-domains in the regions shown in green in **Figure 1A-E**, we were able to engineer exchanges between them in this study. *P. berghei* parasites expressing TRAP without an I-domain show the I-domain is essential for motility and invasion. Parasites expressing TRAP with I-domains from other organisms including humans can have nearly intact motility and invasion. However, a poly-specific, cationic integrin αI-domain functions much better than a more specific, anionic integrin αI-domain. Together with altering the charge of the native TRAP adhesin, our results suggest that cationic surface charge around the MIDAS motif can enable polyspecificity and permit TRAP to function as an adhesin with diverse ligands in both vertebrate and arthropod hosts.

**Figure 1.**
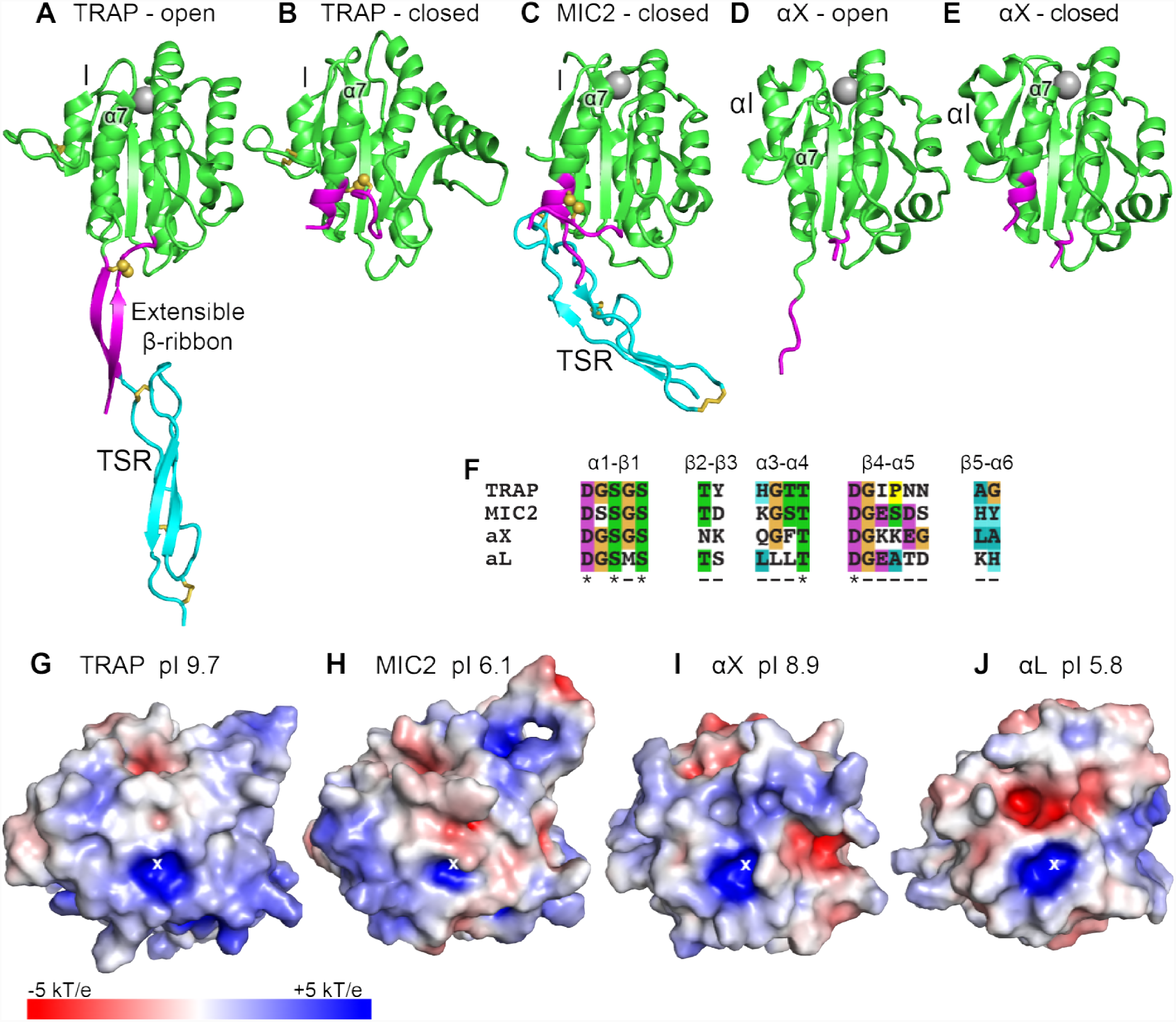
Structural features of TRAP, MIC2, and integrin αI-domains. **A-E)** Cartoon ribbon diagrams. **A**, open TRAP (Protein Databank ID (PDB)) 4HQL; **B**, closed TRAP, PDB 4HQF; **C**, closed MIC2, PDB 4OKR chain B; **D**, open αX αI-domain, PDB 4NEH; **E**, closed αX αI-domain, PDB 5ES4. The portion of the I-domains exchanged between TRAP and other I-domains is shown in green. The TSR domain is shown in cyan. The portion in between the I-domain and TSR in TRAP and MIC2 which includes the extensible β-ribbon is shown in magenta; the comparable regions in integrin αI-domains are also colored magenta and emphasize how the agehelix reshapes similarly to that of apicomplexan I-domains. The TSR domain of closed TRAP **(B)** is not shown because it was disordered in crystals; it likely is positioned similarly in the closed conformation as the TSR domain of closed MIC2 **(C)**. The MIDAS Mg^2+^ ion is shown as a silver sphere and is present in all I-domains. It is not shown in closed TRAP **(B)** because a lattice contact disrupted the conformation around the MIDAS. Disulfide bonds are shown as yellow sticks; for emphasis, the cysteine sulfur atoms of the extensible I-domains are shown in identical orientations after superimposition. **F)** Sequences of the five loops that surround the MIDAS metal ion in the I-domains exchanged here. Residues that are surface exposed and may contact ligands are underlined. Residues that coordinate the MIDAS metal ion directly or indirectly through a water molecule are asterisked. **G—J)** Electrostatic surfaces around the MIDAS metal ion of I-domains in the open conformation. Structures are of G, *P. berghei* TRAP modeled on *P. vivax*, PDB 4HQL; H, MIC2, modeled on closed MIC2 (PDB 4OKR chain B) and open TRAP (PDB 4HQL); I, integrin αX, PDB 4NEH; and J, integrin αL, PDB 1MQ8.

## Results

### The I-domain of TRAP is crucial for *Plasmodium* transmission

We first generated a parasite line expressing TRAP that lacked the I-domain but was otherwise intact (*trapΔI*) (**Figure 2A, Figure S1**). In three different experiments in which mosquitoes fed on infected mice, no *trapΔI* as well as *trap(-)*sporozoites could be observed within mosquito salivary glands while an average of ∼10,000 sporozoites per mosquito was counted for wild-type (**Figure 2B, Table 1**). These results show that *trapΔI* sporozoites are severely impaired in salivary gland invasion. To determine whether mutant parasites retained the ability to migrate steadily on microscope slides; i.e. to glide, sporozoites were isolated from hemolymph and activated by addition of 3% bovine serum albumin (BSA). Gliding motility is a form of circular movement typically displayed by sporozoites (Vanderberg 1974). In this study we defined sporozoites as gliding and productively motile if they were able to complete at least one circle within 3 or 5 minutes, depending on the experiment. Sporozoites exhibiting other types of motion were classified as unproductively motile, while sporozoites that were attached but were not moving or were not attached were classified as non-motile (**Figure S2**) (Münter et al. 2009). Assays with hemolymph (HL) sporozoites revealed that ∼19% of wild-type sporozoites productively moved in a circular fashion while no productive motility was observed for *trapΔI* and *trap(-)* sporozoites (**Figure 2C**). These results showed that the I-domain is required for productive motility. In addition, the infectivity of sporozoites was tested by exposing naive mice to infected mosquitoes as well as by intravenously injecting 10,000 hemolymph (HL) sporozoites. Upon infection with wild-type, the first blood stage parasites were visible after ∼3 days independent of the infection route; in contrast, no infections could be observed for *trapΔI* (**Figure S3, Table 2**) as previously reported for *trap(-)* (Sultan et al. 1997; Sultan et al. 2001). To test if *trapΔI* parasites express TRAP, immunofluorescence assays with antibodies to the TRAP repeat region were performed on isolated midgut sporozoites. TRAP-specific fluorescence was observed in most sporozoites at one end with no difference between *trapΔI* and wild-type sporozoites. This suggests that the mutated TRAP is also localized in the secretory micronemes. In contrast, TRAP-specific fluorescence was completely absent in *trap(-)* sporozoites. TRAP could also be observed on the surface of unpermeabilized *trapΔI* sporozoites indicating that micronemal secretion is not affected in these parasites (**Figure 2D**).

**Table 1.**
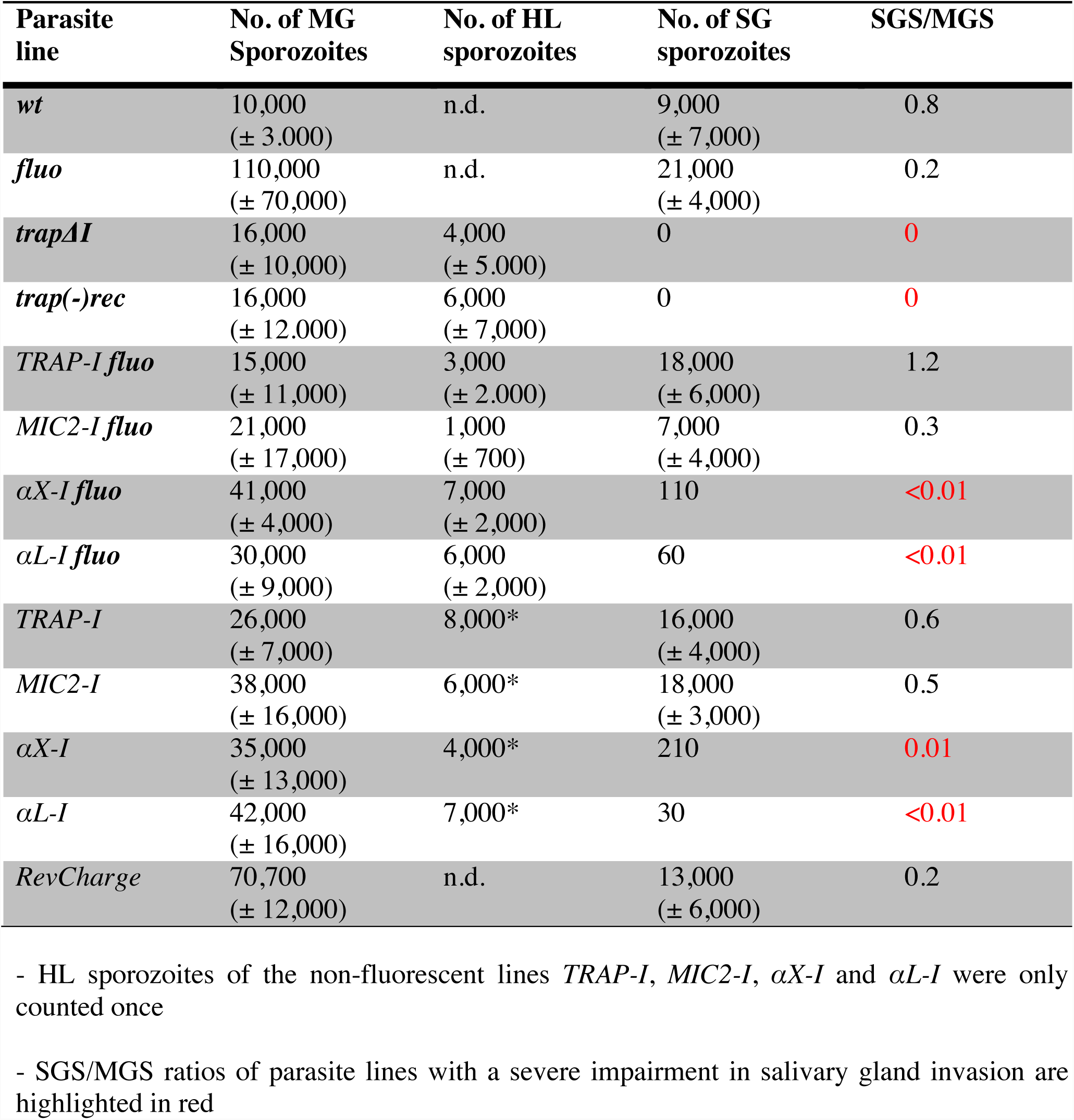
Absolute sporozoite numbers in midgut (MG), hemolymph (HL) and salivary glands (SG) of all analyzed parasite strains. Sporozoites in the midgut, hemolymph and the salivary glands of infected mosquitoes were counted between day 14 and day 24 post infection of each feeding experiment. Shown is the mean ± SD of all countings performed per line. Note that only mosquitoes infected with fluorescent parasites were pre-selected, hence sporozoite numbers per infected mosquito for non-fluorescent lines might be higher. n.d. – not determined.

**Table 2.** Determination of transmission efficacy *in vivo*. Transmission efficacy of the generated parasite lines *trapΔI, TRAP-I*, *MIC2-I*, *αX-I*, *αL-I* and *RevCharge* in comparison to the reference lines *fluo* and wild-type (*wt*). The prepatency is determined as the time between sporozoite infection and the first observance of blood stages and is given as the mean of all positive mice of the respective experiment(s). C57BL/6 mice were either injected intravenously (i.v.) with 10,000 salivary gland sporozoites (SGS) or 10,000 hemolymph sporozoites (HLS) or exposed to 10 mosquitoes that received an infected blood meal. Note that mosquitoes infected with fluorescent parasite lines were pre-selected for parasites by fluorescence in the midgut. Mosquitoes infected with non-fluorescent parasites were not pre-selected but dissected afterwards to ensure that all mice were bitten by at least one infected mosquito. For strains with strongly decreased salivary gland invasion capacity (*aX-I* and *aL-I*) no SGS but 25,000 HLS were injected.

| Parasite line | Route of Inoculation | Mice<sup>1)</sup> | Prepatency (days) |
| --- | --- | --- | --- |
| *fluo* | by mosquito bite<sup>2)</sup> | 4/4 | 3.0 |
| *fluo* | by mosquito bite<sup>3)</sup> | 4/4 | 3.0 |
| *fluo* | 10,000 SGS (i.v.) | 4/4 | 3.5 |
| *TRAP-I fluo* | by mosquito bite<sup>2)</sup> | 4/4 | 3.0 |
| *TRAP-I fluo* | 10,000 HLS (i.v.) | 4/4 | 4.0 |
| *TRAP-I fluo* | 10,000 SGS (i.v.) | 4/4 | 3.0 |
| *MIC2-I fluo* | by mosquito bite<sup>2)</sup> | 4/4 | 3.0 |
| *MIC2-I fluo* | 10,000 HLS (i.v.) | 4/4 | 3.8 |
| *MIC2-I fluo* | 10,000 SGS (i.v.) | 8/8 | 3.1 |
| *αX-I fluo* | by mosquito bite<sup>2)</sup> | 0/4 | /* |
| *αX-I fluo* | 10,000 HLS (i.v.) | 1/4 | 6.0 |
| *αX-I fluo* | 25,000 HLS (i.v.) | 2/4 | 6.0 |
| *αL-I fluo* | by mosquito bite<sup>2)</sup> | 0/4 | /* |
| *αL-I fluo* | 10,000 HLS (i.v.) | 0/4 | /* |
| ***αL-I fluo*** | 25,000 HLS<br>(i.v.) | 1/4 | 5.0 |
| ***wt*** | by mosquito<br>bite<sup>3)</sup> | 8/8 | 3.1 |
| ***wt*** | 10,000 HLS<br>(i.v.) | 4/4 | 3.0 |
| ***wt*** | 10,000 SGS<br>(i.v.) | 4/4 | 3.25 |
| ***trapΔI*** | by mosquito<br>bite<sup>3)</sup> | 0/4 | /* |
| ***trapΔI*** | 10,000 HLS<br>(i.v.) | 0/4 | /* |
| ***TRAP-I*** | 10,000 HLS<br>(i.v.) | 4/4 | 3.0 |
| ***MIC2-I*** | 10,000 HLS<br>(i.v.) | 4/4 | 4.0 |
| ***αX-I*** | 10,000 HLS<br>(i.v.) | 3/4 | 5.3 |
| ***αL-I*** | 10,000 HLS<br>(i.v.) | 0/4 | /* |
| ***RevCharge*** | by mosquito<br>bite<sup>3)</sup> | 2/8 | 5.5 |
| ***RevCharge*** | 10,000 SGS<br>(i.v.) | 9/9 | 3.9 |

\*mice did not become positive within ≥10 days post infection.
<sup>1)</sup> Infected mice / inoculated (exposed) mice
<sup>2)</sup> mosquitoes were pre-selected for parasites
<sup>3)</sup> mosquitoes were not pre-selected for parasites
- Results obtained from non-fluorescent lines are shaded in grey while experiments performed with fluorescent lines are shaded in blue

**Figure 2.**
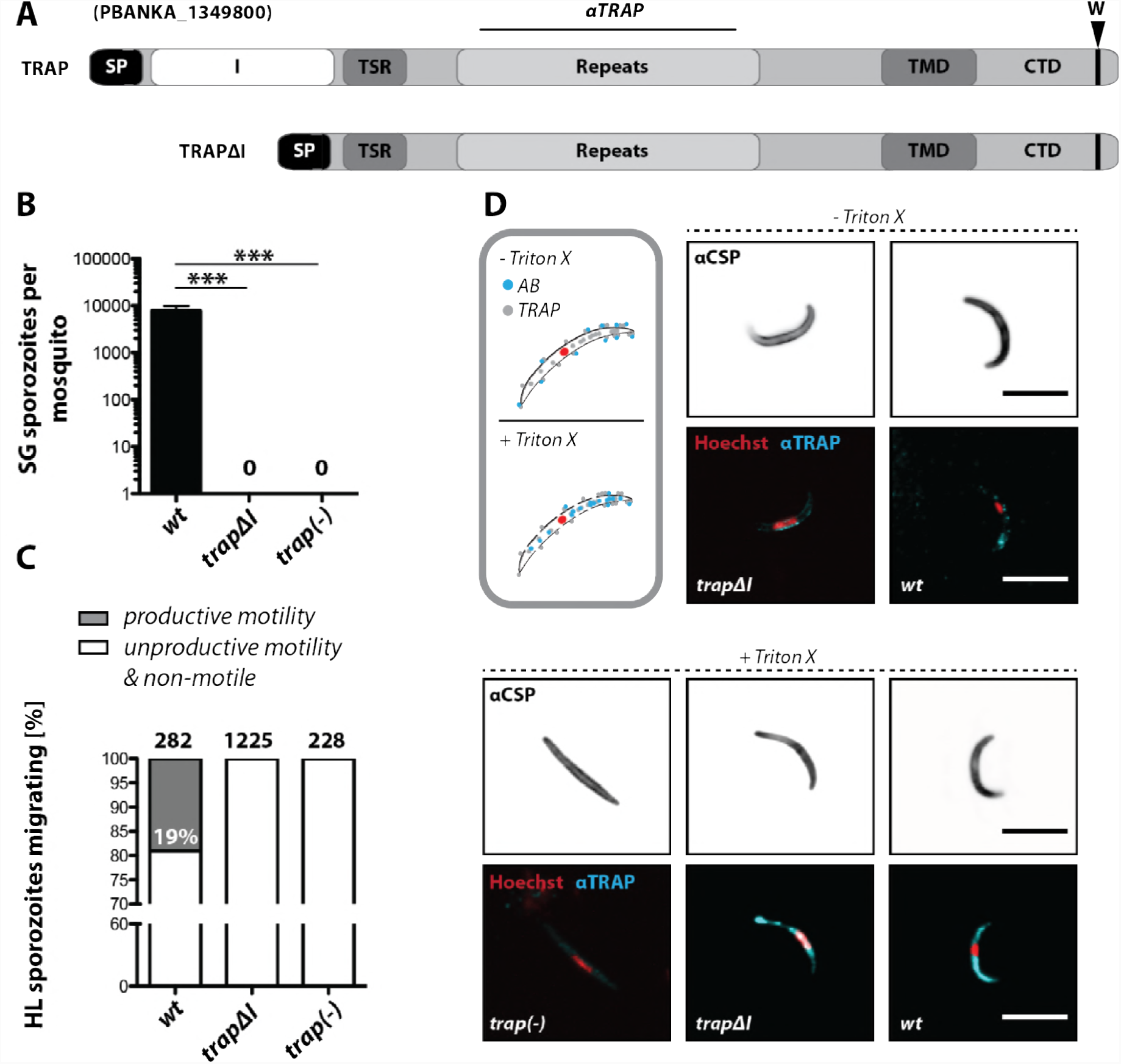
The I-domain of TRAP is essential for salivary gland invasion and gliding motility of sporozoites. **A)**Domain architecture of full-length TRAP and the mutant TRAPΔI lacking the I-domain. TRAP contains a signal peptide (SP) and a conserved penultimate tryptophan (W) as well as a transmembrane domain (TMD) and a cytoplasmic tail domain (CTD). The I-domain and the thrombospondin type-I repeat (TSR) are shown in white and dark grey, respectively. **B)**Sporozoite numbers in the salivary glands of mosquitoes infected with *trapΔI*, *trap(-)* and wild-type (*wt*) 14-22 days post infection. Shown is the mean ± SEM of at least seven counts of three different feeding experiments. ***p<0.0001 one-way-ANOVA (Kruskal-Wallis-test). **C)**Motility of hemolymph sporozoites isolated 13-16 days post infection. Sporozoites moving in at least one full circle within five minutes were considered to be productively moving while all sporozoites that behaved differently were classified as unproductively moving / non-motile. The number of analyzed sporozoites is indicated above each bar. **D)** Immunofluorescence of permeabilized and unpermeabilized midgut sporozoites of *trapΔI* in comparison to wild-type (*wt*) and *trap(-)*. Midgut-derived sporozoites were incubated with αTRAP antibodies recognizing the repeat region. Sporozoites were additionally stained against the surface protein CSP and DNA (Hoechst). Note that intracellular TRAP is stained after permeabilization as indicated in the schematic (grey box). Scale bars: 10 *µ*m.

### Structurally conserved I-domains of other species partially rescue salivary gland invasion

We tested whether the lack of productive motility and infectivity as well as the severely impaired salivary gland invasion rate in *trapΔI* parasites could be complemented by inserting in TRAP an I-domain from a foreign species. We selected structurally characterized I-domains of MIC2 from *Toxoplasma gondii* and the I-domains of the human integrin α-subunits αX (CD11c) and αL (CD11a) (**Figure 1**). Sequence identity among I-domains is 36% between αL and αX, 18% between the integrins and *P. berghei* TRAP, and 28% between TRAP and MIC2 (**Data S1**). Both TRAP and αX are basic, with pI values of 9.7 and 8.9, while MIC2 and αL are acidic, with pI values of 6.1 and 5.8, respectively (Song et al. 2012; Song & Springer 2014). Furthermore, the αX I-domain is poly-specific as shown by binding to multiple glycoproteins and proteolytic fragments as well as heparin (Vorup-Jensen et al. 2005; Vorup-Jensen et al. 2007), while the αL I-domain is highly specific for the ligand intercellular adhesion molecule (ICAM-1) and its homologues ICAM-2, ICAM-3, and ICAM-5 (Grakoui et al. 1999).

The *fluo* line, with eGFP constitutively expressed in all parasite stages and mCherry specifically expressed in sporozoites, was created to simplify analysis of TRAP I-domain replacements throughout the life cycle (**Figure S4**). *Fluo* sporozoites express both fluorescent markers and have salivary gland invasion rates and gliding velocities similar to wild-type (**Figure S5**). Whether sporozoites were administered by infected mosquito bite or injected intravenously, infectivity of the *fluo* line and wild-type were comparable. In line with these results, growth of blood stages and lethality of *fluo* and wild-type parasites were similar (**Figure S6**).

Parasite lines expressing either wild type TRAP (TRAP-I) or TRAP with foreign I-domains (MIC2-I, αX-I or αL-I) replacing the *P. berghei* TRAP I-domain (**Data S1**) were generated with both *fluo* and *non-fluo* (*trap(-)rec)* parasite lines by homologous recombination (**Figures 3A and S7**). We verified the expression of TRAP in *TRAP-I* and *MIC2-I* midgut and salivary gland sporozoites as well as in *αX-I* and *αL-I* midgut sporozoites by immunofluorescence. A distinct TRAP-specific signal was present in all lines often concentrating at the apical tip of the sporozoite (**Figures 3B)**. Sporozoites were isolated from midgut, hemolymph, or salivary glands and quantitated in a hemocytometer or imaged by fluorescence in salivary glands (SG) ≥17 days post infection. TRAP-I *fluo* and *non-fluo* lines gave salivary gland invasion rates that were indistinguishable from *wt* (**Table 1**). Remarkably, as shown both by live fluorescent imaging of *fluo* sporozoites in the SG and quantitation of *fluo* and *non-fluo* lines, the numbers of SG *MIC2-I* and *TRAP-I* sporozoites were comparable at 16,000-18,000 and 7,000-18,000 sporozoites/SG, respectively (**Figure 3C, E, F, Table 1**, and **Movie S1**). Also remarkably, even *αX-I* and *αL-I* sporozoites were capable of invading the SG at low rates (**Table 1** and **Figure 3E** and **F**, note the log scale of the y axis). The numbers for *αX-I* ranged from ∼110 (*fluo*) to ∼210 (*non-fluo*) and for *αL-I* from ∼30 (*non-fluo*) to ∼60 (*fluo*) sporozoites per SG (**Figure 3E, F and Table 1**). We were also able to detect *αX-I* although not *αL-I* sporozoites in the SG by live fluorescent imaging (**Figure 3C**). Sporozoite numbers in the hemolymph were somewhat lower for *MIC-I* and *TRAP-I* than *αX-I* and *αL-I* (**Figure 3D and Table 1**), likely reflecting the efficient SG invasion of *MIC-I* and *TRAP-I* sporozoites.

**Figure 3.**
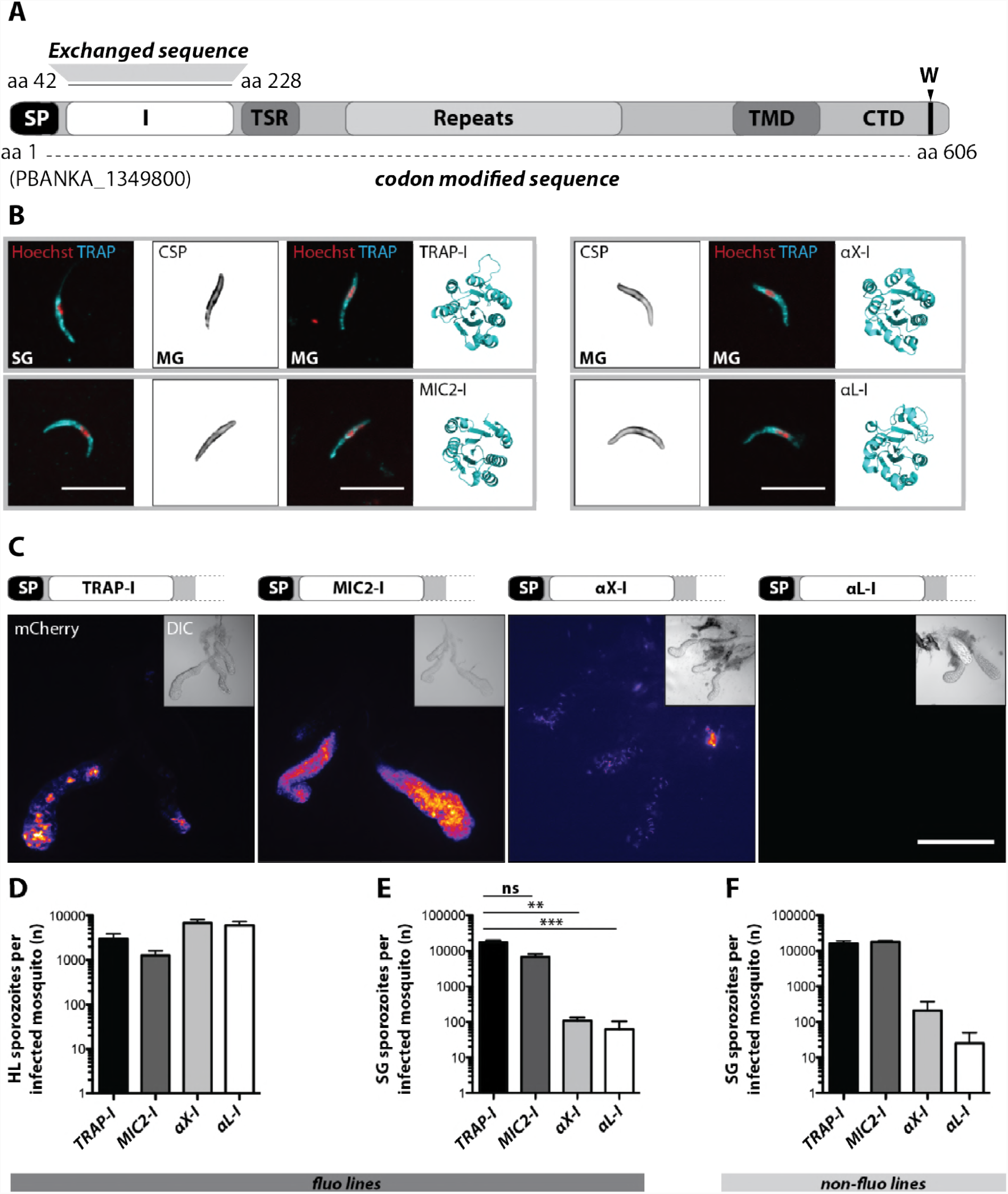
Sporozoites expressing TRAP with I-domains from other species can invade salivary glands. Domain architecture of full-length TRAP (see Figure 2A legend) indicating the exchanged I-domain. **B)** Immunofluorescence assay (IFA) against TRAP and CSP on midgut sporozoites (MG) of *TRAP-I*, *MIC2-I*, *aX-I* and *aL-I* parasite lines. For *TRAP-I* and *MIC2-I* the IFA was also performed with salivary gland sporozoites (SG; on the left). The cartoon ribbon diagram beside each row of images depicts the respective I-domain expressed. IFAs were performed with non-fluorescent parasites (*non-fluo* lines). Scale bar: 10 *µ*m. **C)** Salivary gland colonization of sporozoites expressing the I-domains shown above each image of salivary glands 17-24 days post infection. The fluorescence of mCherry expressing sporozoites is coded for intensity. The small image inset depicts the respective salivary gland in differential interference contrast (DIC). Scale bar: 200 *µ*m. Invasion of the salivary glands by *αX-A* sporozoites is also shown in **Movie S1**. **D and E)** Numbers of hemolymph (HL)-derived (D) and SG-derived (E) sporozoites from mosquitoes infected with the indicated *fluo* line I-domain mutants 13 – 16 days or 17-24 days post infection, respectively. Shown is the mean ± SEM of at least three countings from three independent feeding experiments. **p<0.05 one-way-ANOVA (Kruskal-Wallis-test). **F)** Numbers of salivary gland (SG)-derived sporozoites from mosquitoes infected with *non-fluo* line I-domain mutants. Salivary glands were isolated 17-24 days post infection. Note that mosquitoes infected with non-fluo lines were not preselected for fluorescent parasites before dissection. Graphs show the mean ± SEM of at least two counts per line from a single feeding experiment.

### Divergent I-domains can partially restore gliding motility and infectivity of sporozoites

We next analyzed motility of the parasite lines *in vitro* using the classification scheme shown in **Figure S2**. Gliding assays of hemolymph sporozoites revealed that ∼24% of *TRAP-I* but only ∼4% of *MIC2-I* sporozoites were productively motile **(Figure 4A**). A higher proportion of SG sporozoites were productively motile; ∼53% for *TRAP-I* and ∼15% for *MIC2-I* sporozoites **(Figure 4B**). Among hemolymph sporozoites, ∼1% of *αX-I* sporozoites moved productively; however, none of the >3000 observed *αL-I* sporozoites showed productive movement (**Figure 4A**). *TRAP-I* and *MIC2-I* SG sporozoites that glided continuously for >150 s each had a speed of ∼1.5 *µ*m/s (**Figure 4C**). Furthermore, the trajectories of these continuously gliding sporozoites were similar (**Figure 4D**). Morever, the TRAP-I and MIC2 I-domains supported similarly persistent gliding (**Figure 4E**). Owing to the low numbers of *αX-I* and *αL-I* sporozoites in the SG (**Figure 3**), similar quantitation of sporozoite motility was not possible.

**Figure 4.**
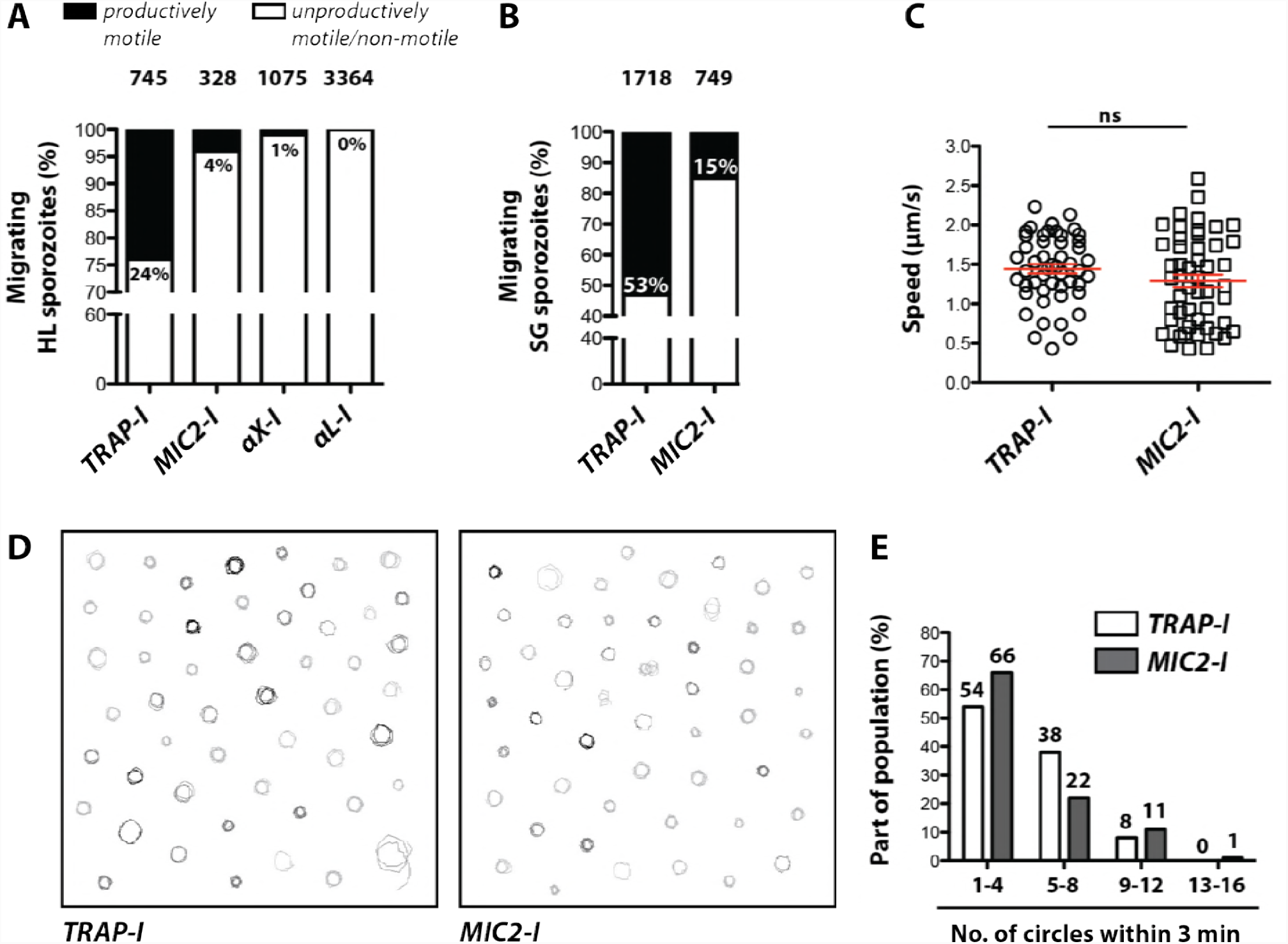
Gliding of I-domain mutant sporozoites. **A, B)** Ratio of productively and unproductively moving / non-motile hemolymph (HL) and salivary gland (SG) (B) sporozoites of the indicated parasite lines. Sporozoites were classified as productively moving if they were able to glide at least one complete circle during a three-minute movie. All sporozoites that behaved differently were classified as unproductively moving / non-motile. Data were generated from three independent experiments per parasite line. The number of analyzed sporozoites is depicted above each column. For further details about movement pattern of sporozoites see Figure S2. Note that SG sporozoites from the lines *αX-I* and *αL-I* could not be analyzed because of the low number of SG sporozoites. Data were generated from three independent experiments per parasite line and the numbers of analyzed sporozoites are depicted above each column. **C)** Average speed of SG sporozoites. Only sporozoites that showed continuous gliding for 150 seconds were analyzed. Shown are the mean speeds of 50 SG sporozoites per line generated from two (*MIC2-I*) and three (*TRAP-I*) independent experiments. Data were tested for significance with an unpaired t-test. Red bars show the mean ± SEM. **D)** Trajectories of the sporozoites tracked in C. **E)** Persistence of gliding *TRAP-I* and *MIC2-I* sporozoites. The graph illustrates the number of circles SG sporozoites of both lines were able to glide during a three-minute movie. Data were generated from two (*MIC2-I*; 107 sporozoites) and three (*TRAP-I*; 111 sporozoites) independent experiments.

To probe the infectivity of mutant sporozoites, mice were exposed to infected mosquitoes or inoculated by intravenous injection of sporozoites. Exposure of mice to parasitized mosquitoes effectively transmitted *TRAP-I* and *MIC2-I* sporozoites, with an infection rate of 100% with a similar prepatent period (time until an infection could be detected in the blood) of three days (**Table 2**). In contrast, no transmission by mosquito bite could be observed for *αX-I* and *αL-I* sporozoites (**Figure S8, Table 2**). Intravenous injection of 10,000 *TRAP-I* or *MIC2-I* SG sporozoites also infected 100% of mice with a prepatency of three days; however, the numbers of *αX-I* and *αL-I* SG sporozoites were too low for similar tests (**Table 2, Figure S8**). We therefore turned to HL sporozoites to compare all four parasite lines for their ability to mediate infection. Injections with 10,000 *TRAP-I* and *MIC2-I* HL sporozoites were able to infect all mice, with slightly longer prepatency of three to four days compared to SG sporozoites (**Figure S9, Table 2**). Interestingly, 50% (4/8) of mice injected with 10,000 *αX-I* HL sporozoites became blood stage patent after a delayed prepatency of >5 days while no infection was observed for the same number of *αL-I* HL sporozoites (0/8) (**Figure S9, Table 2**). In a fourth experiment with 25,000 HL sporozoites, 2 of 4 mice injected with *αX-I* sporozoites became blood stage patent and additionally, 1 of 4 of mice injected with *αL-I* sporozoites became infected (**Figure S9, Table 2**). To exclude spurious results from contamination with other parasite lines, blood-stage transmitted parasites were propagated in mice and analyzed via PCR for the correct genotype (**Figure S10**). Additionally, the TRAP locus of these parasites was sequenced to ensure that the correct I-domain was present. In all tested cases, the expected *αX-I* and *αL-I* genotype was confirmed. Thus, *TRAP-I* and *MIC2-I* sporozoites are comparably infectious to mice. *αX-I* and *αL-I* sporozoites are moderately or severely deficient in infectivity, respectively, but are nonetheless infectious.

### Sporozoites expressing the I-domain of *Tg*MIC2 are impaired in hepatocyte invasion but do not show altered host cell tropism

While *Plasmodium* sporozoites can infect different types of cells they have a strong tropism for the liver. In contrast, *Toxoplasma gondii* can infect any nucleated cell from a warm-blooded animal (Boothroyd 2009). Hence, we tested whether an exchange of the I-domain affected tissue tropism of sporozoites in the vertebrate host. In a first step we tested the capacity of *TRAP-I* and *MIC2-I* SG sporozoites to invade HepG2 cells. After exposure for 1.5 hours to confluently grown HepG2 cells, 2-fold fewer MIC2-I sporozoites than TRAP-I sporozoites were intracellular (**Figure 5A**). We also examined the ability following infection of HepG2 cells to develop into the extra-erythrocytic form of the parasite characteristic for the infection of the liver. At 48 hours after infection, 5-fold fewer *MIC2-I* sporozoites had developed into liver stage parasites compared to *TRAP-I* sporozoites (**Figure 5B**). However, the area within cells occupied by growing liver stage parasites after 48 hours was comparable for *TRAP-I* and *MIC2-I* implying that an exchange of the I-domain affects parasite invasion but not intracellular development (**Figure 5C, D**).

**Figure 5.**
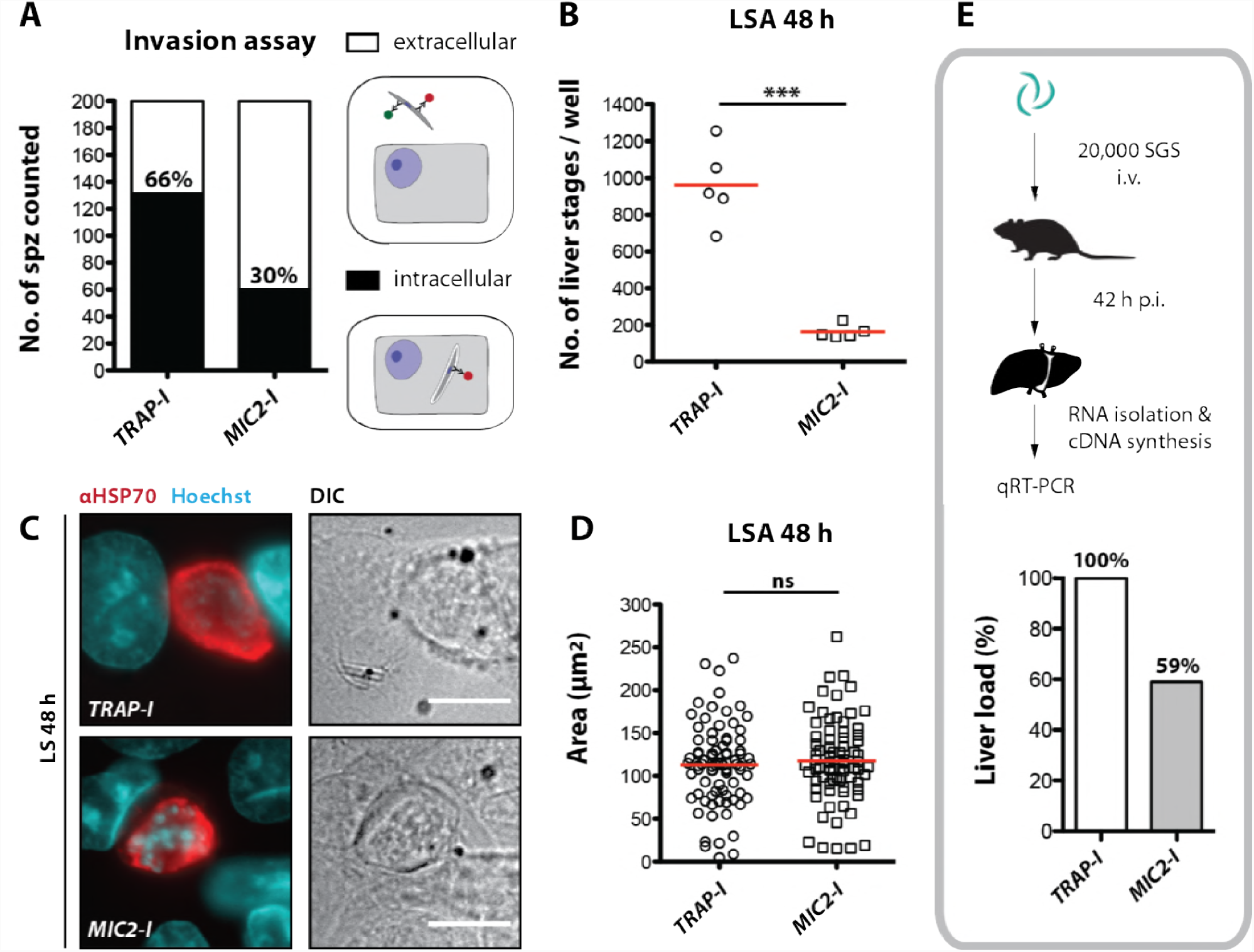
*MIC2-I* sporozoites are impaired in hepatocyte invasion but not in liver stage development. **A)**Invasion assay. Confluent HepG2 monolayers exposed to sporozoites for 1.5 hours were fixed and stained with CSP antibodies before and after permeabilization with methanol; anti-IgG secondary antibodies conjugated to different fluorochromes were used before and after permeabilization. The illustration on the right depicts the staining of intracellular and extracellular sporozoites. **B)** Number of liver stages after challenge with 10,000 SG sporozoites. Experiments were performed with sporozoites obtained from two different feeding experiments with 2-3 technical replicates per parasite line and experiment. The red line indicates the median. *** depicts a p-value of 0.005; unpaired t test. **C,D)** Liver stage development of TRAP-I and MIC2-I parasites. **C)** Immunofluorescence images of *TRAP-I* and *MIC2-I* liver stages 48 hours post infection. The expression of *Pb*HSP70 is shown in red while DNA is shown in cyan. Scale bars: 10 *µ*m. **D)** Area of liver stages 48 hours post infection. The red line shows the median. Data were tested for significance with an unpaired t test. **E)** Liver load in mice challenged with *MIC2-I* as % of load with the control *TRAP-I*. Shown is the relative expression of parasite-specific 18s rRNA determined by qRT-PCR and normalized to mouse-specific GAPDH. Data display the mean of three measurements generated from pooled cDNA of four challenged mice per parasite strain.

We further tested whether *in vivo* tropism of sporozoites was altered by the MIC2 I-domain. Mice were infected by intravenous injection of 20,000 *TRAP-I* and *MIC2-I* SG sporozoites and after 42 hours liver, spleen, lung and a part of the small intestine were harvested. Parasite load was measured by quantitative RT-PCR. Liver tropism of both *TRAP-I* and *MIC2-I* SG sporozoites was pronounced, with ∼32-fold more parasites localizing to the liver than to any other organ (**Figure 5E, Figure S11**). However, the liver burden of *MIC2-I* parasites was reduced ∼40% relative to *TRAP-I* parasites, reflecting the similar decrease observed during *in vitro* infection experiments.

### Charge reversal mutations of the TRAP I-domain differentially impacts infectivity

To determine whether the cationic charge of the I-domain was functionally important, we rendered it anionic with charge reversal mutations. Seven mutations (H56E, H62E, H123E, K164Q, K165D, R195E and K202E) were introduced around the perimeter of the putative ligand binding site on the I-domain, distal from the MIDAS. The predicted pI of the *P. berghei* ANKA strain TRAP I-domain of 9.7 was altered by the mutations to 6.8 (**Figure 6A**). Secretion of TRAP to the surface was not altered in *RevCharge* sporozoites and no difference in protein sub-cellular distribution or expression was observed (**Figure 6B, C**) Salivary gland invasion did not significantly differ (**Figure 6D, Table 1**). Strikingly however, only ∼1% of *RevCharge* SG sporozoites were productively motile in gliding assays compared to ∼53% of *TRAP-I* parasites (**Figure 6E**). The infectivity of *RevCharge* was tested by intravenous injection of 10,000 SG sporozoites into mice in three independent experiments (9 mice in total). While all injected mice became blood stage patent, the prepatency period was delayed for 0.5 - 1 day compared to controls (*wt / TRAP-I / fluo*) (**Figure 6F, Table 2**) corresponding to a decreased infectivity by 50-90%. The decrease in transmission efficacy of *RevCharge* sporozoites by mosquito bite was more marked. Only 2 out of 8 mice in two independent experiments became blood stage positive with a delay in prepatency of >2 days (Figure 6G, **Table 2**) indicating a decreased infectivity by over 99%. These results suggest that the basic charge of the TRAP I-domain is a key determinant of sporozoite infectivity during natural transmission.

**Figure 6.**
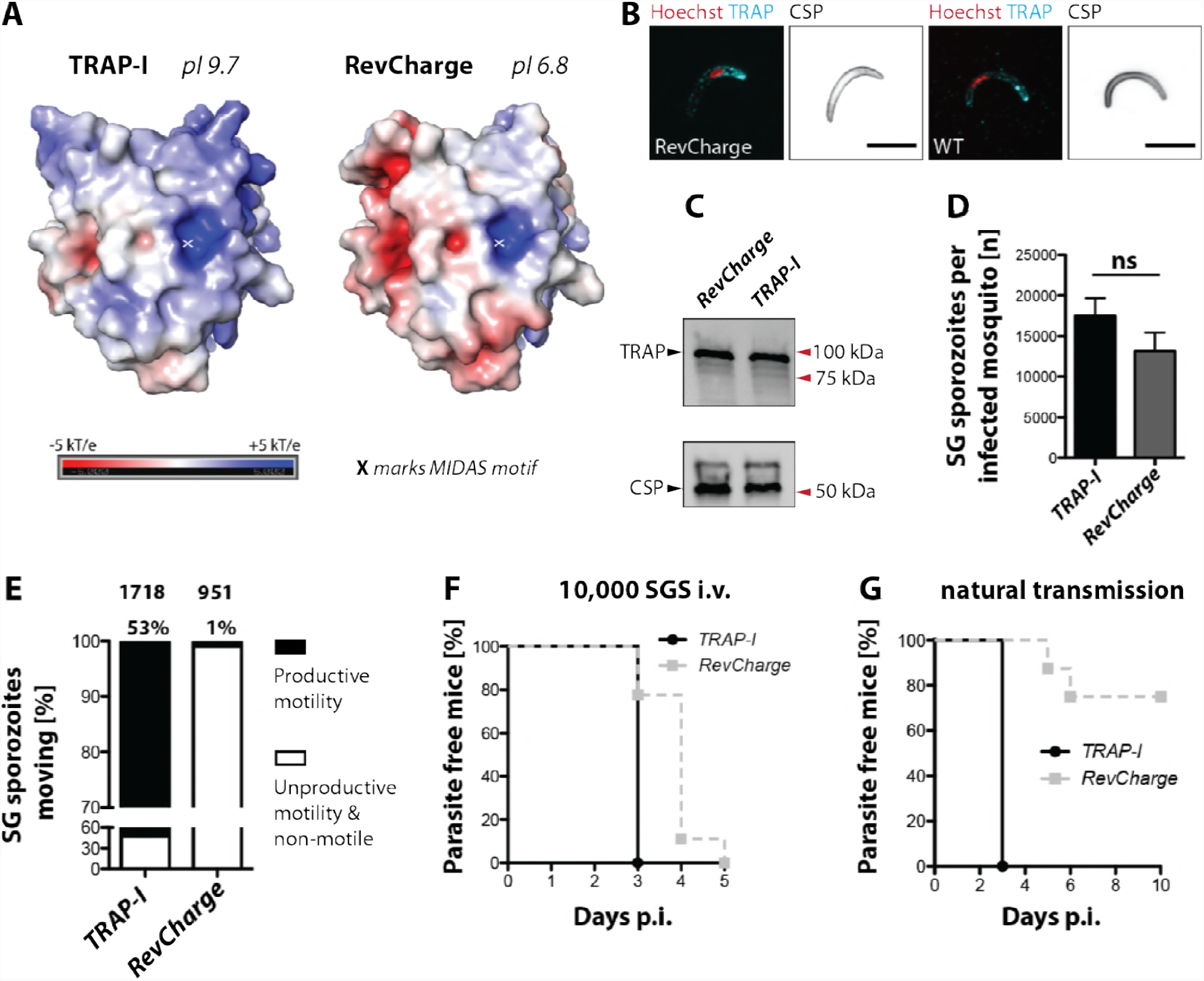
Sporozoites expressing an I-domain with a negatively charged surface show decreased motility and infectivity. **A)** Electrostatic surfaces around the MIDAS metal ion (marked with an asterisk) of I-domains in the open conformation. Structures are of *P. berghei* TRAP (left) and RevCharge (right), respectively, modeled on open *P. vivax*, PDB 4HQL. The color code refers to the electrostatic potential of the surface ranging from −5 to +5 kT/e. **B)** Fluorescence staining on unpermeabilized salivary gland sporozoites of RevCharge and wild-type (*wt*). TRAP is indicated in cyan, DNA in red, and CSP in black. Scale bar: 10 *µ*m. **C)** Western blot on salivary gland (SG) of the control (TRAP-I) and the RevCharge mutant. The circumsporozoite protein (CSP) was used as loading control (shown below). **D)** Sporozoite numbers in salivary glands of mosquitoes infected with *TRAP-I* (*fluo*) and *RevCharge* (*non-fluo*). Shown data represent counts from three independent feeding experiments per parasite line. Data were tested for significance with the Mann-Whitney test. **E)** Movement pattern of *TRAP-I* and *RevCharge* salivary gland (SG) sporozoites. Data were generated from three different feeding experiments. Sporozoites were classified as productively moving if they were able to perform ≥ 1 circle during a three minute movie. All sporozoites that behaved differently were classified as unproductively moving or non-motile. **F,G)** *In vivo* infectivity of *TRAP-I* and *RevCharge* SG sporozoites. C57BL/6 mice were either exposed to infected mosquitoes or injected with 10,000 SG sporozoites/mouse intravenously. **F**) The percentage of parasite-free mice over time after injection of 10,000 SG sporozoites (TRAP-I, n = 4; MIC2-I, n = 9). **G**) The percentage of parasite-free mice over time after exposure to infected mosquitoes (TRAP-I, n = 4; MIC2-I, n = 8).

## Discussion

### Structurally related I-domains partially rescue TRAP function

Previous studies that investigated the function of the thrombospondin related anonymous protein (TRAP) focused on the thrombospondin type I-domain and invariant residues that coordinate the MIDAS metal ion in the I-domain (Wengelnik et al. 1999; Matuschewski et al. 2002). The sidechains of these five invariant I-domain residues coordinate the MIDAS Mg^2+^ ion either directly or indirectly through water molecules. In contrast to the closed conformation, in the open conformation of TRAP and integrin αI-domains, neither of the two Asp residues directly coordinates the MIDAS metal ion. The metal ion is therefore thought to have high propensity to bind an acidic residue in the ligand. The MIDAS residues and their bound waters occupy five of the six coordination positions around the MIDAS Mg^2+^ ion. The remaining, sixth coordination position is occupied when a critical Asp or Glu sidechain in the ligand binds through a carboxyl oxygen to the MIDAS Mg^2+^ ion (Liddington 2014). Mutation of single MIDAS residues or removal of the Mg^2+^ ion by chelation abolishes ligand binding by integrins (Michishita et al. 1993; Kern et al. 1994; Kamata et al. 1995). Similarly, mutation of the MIDAS motif of TRAP severely impairs salivary gland invasion and infectivity of the vertebrate host while gliding motility is decreased in salivary gland but not hemolymph sporozoites (Matuschewski et al. 2002). In contrast, deletion of TRAP completely abrogates salivary gland invasion, infectivity and productive motility (Sultan et al. 1997; Münter et al. 2009) and severely affects substrate adhesion (Münter et al. 2009; Hegge et al. 2010).

To elucidate how TRAP functions in motility we generated the parasite line *trapΔI* expressing TRAP without its I-domain. Interestingly, this line was completely unable to invade salivary glands or infect mice. Furthermore, we observed no productive movement in *trapΔI* sporozoites, similar to *trap(-)* sporozoites. The observation that like *trap(-)* sporozoites (Münter et al. 2009), some *trapΔI* sporozoites could still move in a back-and-forth manner over a single adhesion site, but failed to form more adhesion sites necessary for motility, shows that the I-domain is also important for general adhesion preceding productive motility.

Surprisingly, our study showed that the replacement of TRAP’s native I-domain with the I-domain of MIC2 or αX still enabled sporozoites to perform productive motility *in vitro*, to invade salivary glands, and to infect mice. The MIC2 I-domain gave SG invasion and infection comparable to wt. Sporozoites with αX or αL I-domains replacing the TRAP I-domain showed ∼100-fold reduction in salivary gland invasion and hemolymph sporozoites showed decreased infectivity, with the decrease greater for αL than the αX. The integrin I-domains may have been less effective than apicomplexan I-domains either because of differences in ligand recognition, differences in energies between closed and open states, or the greater structural and sequence differences between the two portions of the I-domain segments joined together. Structures of TRAP have shown that it can assume both open and closed conformations (Song et al. 2012). Integrin I-domains mediate cell migration and stabilizing them in either their open, high-affinity or closed, low-affinity states blocks cell migration. This finding suggests that integrins must be able to transition between open and closed conformations to mediate cell migration, presumably at sites of adhesion and detachment, respectively. The affinity for ligand and the relative free energies of three conformational states tune integrins to be activated by force that is applied by the actin cytoskeleton, resisted by ligand outside the cell, and experienced by the integrin as a tensile force that stabilizes extension (Li & Springer 2017). While integrins are heterodimers with three conformational states, TRAP is apparently a monomer with two conformational states, and its I-domain may be tuned differently in its response to applied force and the difference in energy between its closed and open states. Differences in tuning may in part explain the lesser efficacy of the integrin αX and αL I-domains compared to the apicomplexan MIC2 I-domain. Furthermore, I-domain sequence identity of 18% between integrins and TRAP is from a structural point of view markedly less than the 28% between TRAP and MIC2. Furthermore, TRAP and MIC2 have disulfide bonds between the N and C-terminal portions of their I-domains, while integrins do not. Incompatibilities between the major portion of the I-domain and the N and C-terminal portions of TRAP into which it was inserted thus are larger for the integrin I-domains than the MIC2 I-domain. Thus, greater sequence, structural, and disulfide bond similarities to TRAP may also have contributed to the much greater efficacy of MIC2 than the integrin I-domains in mediating sporozoite emigration to salivary glands and infection of mammalian hosts. Considering these limitations, it is surprising how well the replacements with the apicomplexan and integrin I-domain functioned in both mediating gliding motility, invasion of salivary glands, and infection of mammalian hosts.

### Structural and charge-dependent interactions ensure TRAP function

As TRAP, MIC2, and the integrins αXβ2 and αLβ2 mediate adhesion and migration in completely different hosts and tissues, the underlying features required for gliding motility, invasion, and infection must be highly conserved in αI-domains. One essential I-domain feature may be the MIDAS motif with its positively charged metal ion and transition between low and high affinity conformations. As integrin αXβ2 is poly-specific while αLβ2 is specific, and αXβ2 like TRAP is basic, we hypothesized that the αX αI-domain would more efficiently replace TRAP I-domain function than αL as indeed we found. As the vast majority of cell surface and extracellular proteins are acidic, basic adhesins can in part bind ligands through charge complementarity. The surface charge of the tested I-domains varies remarkably ranging from 5.8 (αL-I) to 9.7 (TRAP-I). Curiously the I-domain of MIC2 displayed the best results despite its negatively charged (acidic) surface (Song & Springer 2014). Charge alone thus cannot explain the differences among the replacement mutants, and other features including the greater sequence and structural similarity between the apicomplexan I-domains discussed in the preceding paragraph may be important in the efficacy of the MIC2 I-domain. However, altering the charge around the MIDAS motif of TRAP-I severely impaired gliding motility and infectivity of sporozoites to the mammalian host showing that charge-based interactions are important. Interestingly, the RevCharge mutations did not affect invasion of salivary glands, hence revealing an intriguing decoupling of salivary gland infection with infection of the mammalian host in TRAP function. This suggests a differential need for different parts of the I-domain for mosquito salivary gland invasion as well as crossing of the dermis. Intriguingly, MIC2-I is more negatively charged than RevCharge, and also exhibited a decrease in motility but not in infectivity, suggesting that overall charge may be important in motility, but is not sufficient to explain accumulation in salivary gland or liver infections. Taken together our results suggest that both structural and charge-dependent interactions act in concert to ensure efficient binding by TRAP to its ligands.

### TRAP is a poly-specific receptor interacting with a broad range of ligands

Since discovery of TRAP three decades ago (Robson et al. 1988), identification of its ligands has been of great interest. TRAP-ligand interactions have been hypothesized to explain tropism of sporozoites for specific invasion of salivary glands of the mosquito and the liver of the vertebrate host. Proteoglycans as well as liver-specific proteins like Fetuin-A (Robson 1995; Pradel et al. 2002; Jethwaney et al. 2005) have been described to function as putative ligands for TRAP on hepatocytes. In line with these observations, TRAP I-domain MIDAS motif mutations were shown to drastically decrease the capacity to invade the salivary glands and to infect hepatocytes, leading to a model that suggested recognition of host cells by TRAP binding to glycosaminoglycans in combination with unknown ligands (Matuschewski et al. 2002). Recently it was also found that *Pf*TRAP interacts *in vitro* specifically with integrins containing the αV subunit (Dundas et al. 2018). Careful analysis, however, revealed only a minor role for this interaction during migration in the skin. Similarly, a salivary gland specific protein named Saglin was shown to bind TRAP and to be important for sporozoite entry into salivary glands (Ghosh et al. 2009). Interestingly, we showed that replacement of TRAP’s native I-domain with the I-domain of MIC2 from *Toxoplasma gondii* largely restored salivary gland invasion and infectivity, while affecting motility on glass. This complementation is of particular interest because *Toxoplasma gondii* is a very promiscuous parasite that can infect nearly any nucleated cell of warm-blooded animals while its life cycle does not involve passage through an arthropod vector. These observations make it rather unlikely that TRAP recognizes specific cell surface receptors that are exclusively expressed in the salivary glands or on hepatocytes. It might still be possible that other regions of TRAP’s extracellular domain act in concert with its native I-domain to define host cell specificity. However, it was shown before that shortening of TRAP by deletion of the juxtamembrane domain (JMD, region in between transmembrane domain and repeats of TRAP) does not affect TRAP function (Ejigiri et al. 2012). Mutation of the TSR was reported to only have a slight negative effect on salivary gland numbers and infectivity while leaving life cycle progression unaffected (Matuschewski et al. 2002). The only extracellular region not investigated by genetic approaches so far is the repeat region located in between the TSR and the JMD. This region varies hugely in composition and number between different *Plasmodium* species even if they infect the same host (highly repetitive in *P. falciparum* and *P. malariae*, non-repetitive in *P. vivax* and *P. ovale*), which makes a specific role in host cell recognition unlikely, especially as *P. vivax* TRAP can be expressed in place of *P. berghei* TRAP without loss of function (Bauza et al. 2014). In addition to these findings new sequencing data revealed that also bird-infecting malaria species like *P. gallinaceum* and *P. relictum* express TRAP while sporozoites of these species develop in fibroblasts of the skin and are not targeted to the liver of the infected bird (Böhme et al. 2018).

### Cooperation of TRAP with other receptors could define host cell specificity

Putting our observations in perspective we suggest that TRAP acts as a universal adhesin using two independent mechanisms. First it mediates adhesion during gliding motility by charge-dependent interactions of its I-domain with a broad range of substrates. By this mechanism the sporozoite attaches to the substrate whilst transient polyspecific interactions enable fast forward movement. Second, TRAP assists specifically in host cell attachment in a MIDAS-dependent manner that might be required to initiate invasion. Both functions could mesh in a two-step process providing transient adhesion to substrates and cells. However, our exchange studies with other I-domains suggest that MIDAS-dependent cell attachment is polyspecific and does not define host cell tropism. This implies that either other adhesion receptors or chemoattractants target sporozoites to the correct organs to enable TRAP-mediated entry. The 6-cysteine domain protein p36 has been described to be crucial for liver infection of *Plasmodium spp.* by mediating interactions with the surface receptors EphA2, CD81 and the scavenger receptor BI (Kaushansky et al. 2015; Manzoni et al. 2017). Also, the circumsporozoite protein CSP was shown to play a key role in the colonization of salivary glands and efficient liver cell entry (Coppi et al. 2011). Alternatively, sporozoites might have receptors that sense substrate stiffness or sense molecules that diffuse from mosquito salivary glands or host cells and that activate the actin cytoskeleton machinery that couples to TRAP. Prokaryotes and eukaryotes have evolved distinctive chemotaxis receptors and downstream signaling pathways. Chemotaxis in sporozoites has been discussed, but remains undefined at the molecular level (Muthinja et al. 2017). Chemoattractants have the advantage that they can drive directional migration through multiple layers of distinctive cell types. Vertebrate leukocytes have chemoattractant receptors that activate the actin cytoskeleton which couples to I-domain-containing integrins. In turn, the integrins bind to extracellular ligands and provide the traction for cell migration through multiple tissue layers. This system enables leukocyte emigration from the vasculature in highly specific organs or sites of inflammation (Springer & Dustin 2012). It is possible that similar coordination between chemoattractant receptors and TRAP or mechanosensing might drive sporozoite tropism for the salivary gland in mosquitos, for the liver in mammalian hosts, and for skin fibroblasts in avian hosts.

In conclusion we show that I-domains from humans can partially and those from *T. gondii* can fully complement the function of the *Plasmodium berghei* TRAP I-domain during infection. Reversing the charge around the MIDAS allowed us to decouple TRAP function in mosquito salivary gland invasion from its functionality in the mammalian host. This suggests that a complex ligand-binding repertoire is embedded in the TRAP I-domain that awaits full characterization.

## Materials and methods

### Bioinformatic analysis

*Plasmodium* sequences were retrieved from PlasmoDB (http://plasmodb.org/plasmo/, version 26-34) (Aurrecoechea et al. 2009) and multiple sequence alignments were performed with Clustal Omega (http://www.ebi.ac.uk/Tools/msa/clustalo/) (Sievers et al. 2011). To change the codon usage of open reading frames (ORFs) we applied the tool OPTIMIZER (http://genomes.urv.es/OPTIMIZER/) (Puigbò et al. 2007).

### Generation of *trap(-)rec* and *trapΔI* parasites

TRAP knockout (*trap(-)*) parasites were generated with the PlasmoGem (Schwach et al. 2015) vector (PbGEM-107890) using standard protocols (Janse et al. 2006). Isogenic *trap(-)* parasites were subsequently negatively selected with 5-fluorocytosine (1.0 mg/mL in the drinking water) to give rise to selection marker free *trap(-)rec* parasites (Braks et al. 2006). For the generation of *trapΔI* parasites we made use of the Pb238 vector (Deligianni et al. 2011; Klug & Frischknecht 2017). In a first step the *trap* 3’UTR (970 bp) was amplified with the primers P165/P166 and cloned (*BamHI* and *EcoRV*) downstream of the resistance cassette in the Pb238 vector. In a next step the coding sequence of the *trap* gene including the 5’ and 3’ UTR was amplified with the primers P508/P509 and cloned in the pGEM-T-Easy vector giving rise to the plasmid pGEM-TRAPfull. Subsequently the pGEM-TRAPfull plasmid was mutated with the primers P535/P536 and P537/P538 to introduce a restriction site for *NdeI* directly in front of the start codon ATG and a restriction site for *PacI* directly after the stop codon TAA. The mutated sequence was cloned (*SacII* and *EcoRV*) in the Pb238 intermediate vector that contained already the *trap* 3’UTR downstream of the selection marker and the resulting plasmid was named Pb238-TRAP-NdeI/PacI. The designed DNA sequence lacking the coding region of the I-domain was codon modified for *E. coli* K12 and synthesized at GeneArt (Invitrogen). Subsequently the designed sequence was cloned (*NdeI* and *PacI*) in the Pb238-TRAP-NdeI/PacI by replacing the endogenous *trap* gene. Final DNA sequences were digested (*NotI*, PbGEM-107890; *SacII* and *KpnI*, Pb238-TRAPΔI), purified and transfected into wild-type parasites (*wt*) using standard protocols (Janse et al. 2006) (**Figure S1**).

### Generation of the selection marker free *fluo* line

To generate a selection marker free parasite line that is strongly fluorescent in sporozoites to enable pre-sorting of infected mosquitoes and to simplify imaging we made use of the Pb262 vector (Kooij et al. 2005; Klug et al. 2016). This vector contained the *mcherry* gene under control of the *csp* promoter and the 3’UTR of the *dhfs* gene. Furthermore the *yfcu-hdhfr* selection cassette in this vector enables positive and negative selection. In a first step the *dhfr* 3’UTR downstream of the selection cassette was removed by site directed mutagenesis (primer P788 and P691) to prevent loss of the selection marker by homologous recombination with its flanking regions. Transfection of this construct into wild-type parasites and subsequent selection with pyrimethamine gave rise to the “docking line” *CSmCherryMinus*. This line cannot lose the selection cassette by homologous recombination but can be forced to replace the selection cassette under drug pressure with DNA sequences that do not contain a selection marker (Lin et al. 2011). This strategy was used to generate the *fluo* line by transfecting a DNA sequence containing the *egfp* gene under control of the *ef1α* promoter flanked by the same sequences as the selection cassette in *CSmCherryMinus* parasites. This sequence was obtained by digesting (*PvuI*) the Pb262CSeGFPef1aeGFP vector that was subsequently transfected into *CSmCherryMinus* parasites. The transfection gave rise to the *fluo* line constitutively expressing eGFP (driven by the *ef1α* promoter) and highly upregulating mCherry expression (driven by the *csp* promoter) in oocysts, sporozoites and liver stages (**Figure S4**).

### Generation of *TRAP-I*, *MIC2-I*, *αL-I*, *αX-I* and *RevCharge* parasites

To generate parasite lines expressing TRAP with different I-domains, bp 115 to bp 696 (581 bp; I42 to V228; 194 aa in total) of the wild-type *trap* gene (*Plasmodium berghei* ANKA strain) were exchanged with sequences from the micronemal protein 2 (MIC2) of *Toxoplasma gondii* (567 bp, L75 to V263, 189 aa), the integrin CD11c/αX (552 bp, Q150 to I333, 184 aa) and the integrin CD11a/αL (531 bp, V155 to I331, 177 aa) of *Homo sapiens*. Chimeric sequences as well as the coding sequence of the wild-type *trap* gene that served as a control, were codon modified for *E. coli* K12 to prevent incorrect integration events with the downstream part of the *trap* coding sequence and to avoid changes of the codon usage within the open reading frame. This enabled also simple differentiation between wild-type and transgenic parasites by PCR. For the generation of *RevCharge* parasites seven mutations of non-conserved amino acids (H56E, H62E, H123E, K164Q, K165D, R195E and K202E; *P. berghei* ANKA strain) were introduced into the codon modified wild-type *trap* gene. These mutations were expected to shift the surface charge at the apical side of the I-domain from a pI of 9.7 to 6.8 while leaving the MIDAS intact and the structural integrity of the domain unaffected. All designed sequences were synthesized at GeneArt (Invitrogen) and cloned in the Pb238-TRAP-NdeI/PacI vector (*NdeI/PacI*) that was already used to generate the *trapΔI* line. Constructs were digested (*ScaI-HF*) and transfected using standard protocols (Janse et al. 2006). Transfections were performed in the negatively selected TRAP knockout line *trap(-)rec* as well as in the fluorescent background line *fluo* to independently generate fluorescent (*fluo*) and non-fluorescent (*non-fluo*) sets of mutants (**Figure S7**).

### Generation of isogenic parasite populations

Isogenic parasite lines were generated by serial dilution of parental populations obtained from transfections. Per transfection one mouse was infected by intraperitoneal injection of ∼200 *µ*L frozen parasites of the parental population. To increase the number of transfected parasites within infected mice pyrimethamine (0.07 mg/mL) was given within the drinking water 24 hours post injection. Donor mice were bled two to three days post injection once parasitemia reached 0.5 – 1%. Parasites were diluted in phosphate buffered saline (PBS) to 0.7 - 0.8 parasites per 100 *µ*L and the same volume was subsequently injected into 6 – 10 naive mice. Parasites were allowed to grow for 8 – 10 days until parasitemia reached 1.5 – 2%. Blood of infected mice was taken by cardiac puncture (usually 600-800 *µ*L) and used to make parasite stocks (∼200 *µ*L infected blood) and to isolate genomic DNA with the Blood & Tissue Kit (Qiagen).

### Mosquito infection

Naive mice were infected with 100 – 200 *µ*L frozen parasite stocks and parasites allowed to grow for four to five days. Infected mice were either directly fed to mosquitoes or used for a fresh blood transfer of 2×10^7^ parasites by intraperitoneal injection into two to three naive mice. Parasites within recipient mice were allowed to grow for further three to four days, depending on the number of exflagellation events observed. To determine the number of exflagellation events, and subsequently the number of male gametocytes, a drop of tail blood was placed on a microscope slide, covered with a coverslip and incubated for 10 – 12 minutes at room temperature (20-22°C). The number of exflagellation events was counted with a light microscope (Zeiss) and a counting grid by using 40-fold magnification with phase contrast. If at least one exflagellation event per field could be observed mice were fed to mosquitoes. Mosquitoes were starved overnight prior to the feeding to increase the number of biting mosquitoes. Per mosquito cage (approximately 200 - 300 female mosquitoes) two to three mice were used for feeding.

### Isolation of midgut, hemolymph and salivary gland sporozoites

To estimate the number of sporozoites, infected *Anopheles stephensi* mosquitoes were dissected on day 13, 14, 18, and 22 post infection. For the collection of hemolymph sporozoites infected *A. stephensi* mosquitoes were cut with a needle to remove the last segment of the abdomen. Subsequently the thorax was pierced with a finely drawn Pasteur pipette filled with RPMI/PS solution. While gently pressing the pipette the haemocoel cavity was flushed with solution which dripped off the abdomen. The sporozoite solution was collected on a piece of foil and transferred into a plastic reaction tube. To determine the number of midgut sporozoites, the abdomen of infected mosquitoes was dissected with two needles and the midgut was extracted. Isolated midguts were transferred into a plastic reaction tube containing 50 *µ*L RPMI/PS solution. For the isolation of salivary gland sporozoites, the head of infected mosquitoes was gently pulled away with a needle while fixing the mosquito in place with a second needle. Ideally the salivary glands stayed attached to the head and could be easily isolated. Salivary glands were transferred into a plastic reaction tube with 50 *µ*L RPMI/PS. To release sporozoites, pooled midguts and salivary glands were homogenized using a plastic pestle for 2 min. To count the parasites in each sample, 5-10 *µ*L of the sporozoite solution (1:10 dilution) was applied on a hemocytometer. Sporozoites were counted using a light microscope (Zeiss) with 40-fold magnification and phase contrast. For each counting experiment at least 10 mosquitoes were dissected. However, the number was adapted depending on the infection rate of the mosquitoes and the experiment that was performed.

### Animal experiments

To determine the prepatency of parasite lines during sporozoite transmission two different routes of infection were tested. Female C57BL/6 mice were either exposed to infected mosquitoes or infected by intravenous injection of hemolymph (HL) or salivary gland (SG) sporozoites. To infect mice by mosquito bites, mosquitoes infected with fluorescent parasite lines were pre-selected for fluorescence of the abdomen by using a stereomicroscope (SMZ1000, Nikon) with an attached fluorescence unit. Subsequently parasite positive mosquitoes were separated in cups to 10 each and allowed to recover overnight. Approximately six hours prior to the experiment mosquitoes were starved by removing salt and sugar pads. Mice were anaesthetized by intraperitoneal injection of a mixture of ketamine and xylazine (87.5 mg/kg ketamine and 12.5 mg/kg xylazine) and placed with the ventral side on the mosquito cups. Mosquitoes infected 17-24 days prior to the experiment were allowed to feed for approximately 15 min before mice were removed. During this time eyes of mice were treated with Bepanthen cream (Bayer) to prevent dehydration of the cornea. After the experiment mice were allowed to recover and tested for blood stage parasites on a daily basis by evaluation of Giemsa stained blood smears. If non-fluorescent parasite lines were tested infected mosquitoes were not pre-sorted. In these experiments midguts of mosquitoes that had taken a blood meal were dissected after the experiment and the number of midgut sporozoites was counted as described previously. Mice that were bitten by mosquitoes that contained no midgut sporozoites were excluded from the analysis.

For injections HL or SG sporozoites were isolated either 13-16 days (HL) or 17-24 days (SG) post infection. Sporozoite solutions were diluted to the desired concentration with RPMI/PS (either 10,000 or 25,000 sporozoites) and injected intravenously into the tail vein of naive mice. The presence of blood stage parasites was evaluated on a daily basis.

### *In vitro g*liding assay

To analyze speed and movement pattern of sporozoites, *in vitro* gliding assays were performed in glass-bottom 96-well plates (Nunc). Hemolymph and salivary gland sporozoites were obtained by dissecting infected *A. stephensi* mosquitoes. To free the sporozoites from salivary glands (SG), samples were grounded with a pestle. Subsequently SG samples were centrifuged for 3 min at 1,000 rpm (Thermo Fisher Scientific, Biofuge primo) to separate sporozoites from tissue. Afterwards ∼40 *µ*L of the supernatant was transferred into a new 1.5 mL plastic reaction tube and diluted with a variable volume of RPMI/PS depending on the planned number of assays and the sporozoite concentration resulting in a minimum of 50,000 sporozoites per well. For each assay about 50 *µ*L of the sporozoite suspension was mixed with 50 *µ*L RPMI medium containing 6% bovine serum albumin (BSA) to initiate activation. Subsequently sporozoites were allowed to attach to the bottom by centrifugation at 800 rpm for 3 min (Heraeus Multifuge S1). Using fluorescence microscopy (Axiovert 200M) with a 10x objective movies were recorded with one image every three seconds for 3 to 5 min depending on the experiment. Movies were analyzed manually using the Manual Tracking Plugin from ImageJ (Schindelin et al. 2012) to determine speed and trajectories of moving sporozoites. Sporozoites that were able to glide at least one full circle during a 3 min movie were considered to be productively moving while all other sporozoites were classified as non-productively moving (moving less than one circle) or non-motile.

### Live imaging

For the imaging of sporozoites within salivary glands infected mosquitoes were dissected 17-24 days post infection as described previously. Isolated salivary glands were transferred with a needle to a microscope slide containing a drop of Grace’s medium (Gibco) and carefully sealed with a cover slip. Samples were imaged with an Axiovert 200M (Zeiss) using 63x (N.A. 1.3) and 10x (N.A. 0.25) objectives.

### Antibodies

For immunofluorescence assays we made use of antibodies directed against the circumsporozoite protein (CSP) and the thrombospondin related anonymous protein (TRAP). In all assays the anti-CSP antibody mAb 3D11 (Yoshida et al. 1980) was applied as unpurified culture supernatant of the corresponding hybridoma cell line (1:5 diluted for immunofluorescence assays). TRAP antibodies were generated against the peptide AEPAEPAEPAEPAEPAEP by Eurogenetec and the purified antibody was applied as 1:100 dilution in immunofluorescence assays. Antibodies against the same peptide have been shown previously to specifically detect TRAP by immunofluorescence and western blotting (Ejigiri et al. 2012). Secondary antibodies coupled to AlexaFluor 488 or Cy5 (goat anti-mouse or goat anti-rabbit) directed against primary antibodies were obtained from Invitrogen and always used as 1:500 dilution.

### Immunofluorescence assays with sporozoites

To visualize the expression and localization of TRAP in sporozoites, infected salivary glands were dissected as described previously and pooled in plastic reaction tubes containing 50 *µ*L PBS or RPMI/PS. Afterwards salivary glands were mechanically grounded with a plastic pestle to release sporozoites from tissue. Immunofluorescence assays were performed by two different methods either fixing sporozoites in solution or on glass cover slips. To fix the parasites on glass, salivary glands were dissected in RPMI/PS and treated as described. Sporozoite solutions were transferred into 24-well plates containing round cover slips, activated with an equal volume RPMI/PS containing 6% BSA and allowed to glide for approximately 30 min at RT. Subsequently the supernatant was discarded and sporozoites were fixed with 4% PFA (in PBS) for 1 h at RT. Fixed samples were washed three times with PBS for 5 min each. If immunofluorescence was performed on sporozoites in solution, salivary glands were dissected in PBS and treated as described previously. Sporozoite solutions were directly fixed by adding 1 mL of 4% PFA (in PBS) for 1 h at RT. After fixation samples were washed as described for samples fixed on glass while samples in solution had to be additionally pelleted after each step by centrifugation for 3 min at 10,000 rpm (Thermo Fisher Scientific, Biofuge primo). Subsequently sporozoites were blocked (PBS containing 2% BSA) or blocked and permeabilized (PBS containing 2% BSA and 0.5% Triton X-100) over night at 4°C or for 1 h at RT, respectively. Samples were incubated with primary antibody solutions for 1 h at RT in the dark and subsequently washed three times with PBS. After the last washing step, samples were treated with secondary antibody solutions and incubated for 1 h at RT in the dark. Stained samples were washed three times in PBS and the supernatant was discarded. If the immunofluorescence assay was performed in solution, sporozoite pellets were resuspended in 50 μL of remaining PBS, carefully pipetted on microscopy slides and allowed to settle for 10– 15 min at RT. Remaining liquid was removed with a soft tissue and samples were covered with cover slips which had been prepared with 7 μL of mounting medium (ThermoFisher Scientific, ProLong Gold Antifade Reagent). If the immunofluorescence assay was performed on sporozoites that were fixed on glass, cover slips were removed with a forceps, carefully dabbed on a soft tissue and placed on microscopy slides that had been prepared with 7 μL of mounting medium. Samples were allowed to set overnight at RT and kept at 4°C or directly examined. Images were acquired with a spinning disc confocal microscope (Nikon Ti series) with 60-fold magnification (CFI Apo TIRF 60x H; NA 1.49).

### Western blot analysis of TRAP expression

Salivary glands of infected mosquitoes were dissected in RPMI medium (containing 50,000 units/L penicillin and 50 mg/L streptomycin) and smashed with a pestle to release sporozoites. Subsequently sporozoites were pelleted by centrifugation for 10 min at 13,000 rpm (Thermo Fisher Scientific, Biofuge primo). The supernatant was discarded and the pellet was lysed in 60 *µ*L RIPA buffer (50 mM Tris pH 8, 1% NP40, 0.5% sodium dexoycholate, 0.1% SDS, 150 mM NaCl, 2 mM EDTA) for ≥1 h on ice. Lysates were mixed with Laemmli buffer, heated for 10 min at 95°C and centrifuged for 1 min at 13,000 rpm (Thermo Fisher Scientific, Biofuge primo). Samples were separated on precast 4–15% SDS-PAGE gels (Mini Protein TGX Gels, Bio-Rad) and blotted on nitrocellulose membranes with the Trans-Blot Turbo Transfer System (Bio-Rad). Blocking was performed by incubation in PBS containing 0.05% Tween20 and 5% milk powder for 1 h at room temperature. Afterwards the solution was refreshed and antibodies directed against TRAP (rabbit polyclonal antibody, 1:100 diluted) or CSP (mAb 3D11, cell culture supernatant 1:50 diluted) were added. Membranes were washed three times (PBS with 0.05% Tween20) and secondary anti-rabbit antibodies (Immun-Star (GAR)-HRP, Bio-Rad) or anti-mouse antibodies (NXA931, GE Healthcare) conjugated to horse radish peroxidase were applied for 1 h (1:10,000 dilution) at room temperature. Signals were detected using SuperSignal West Pico Chemiluminescent Substrate or SuperSignal West Femto Maximum Sensitivity Substrate (Themo Fisher Scientific).

### Liver stage development assay

Two days prior to the experiment 50,000 HepG2 cells/well were seeded in an 8-well Permanox Lab-Tek chamber slide (Nunc). On day 0 salivary glands were isolated from infected female mosquitoes and collected in RPMI medium within a 1.5 mL reaction tube. Sporozoites were released by mechanically disrupting the salivary glands with a polypropylene pestle. The solution was centrifuged in a table top centrifuge at 1,000 rpm for 3 min at RT and the supernatant was transferred to a new 1.5 mL reaction tube. The salivary gland pellet was resuspended in 100 *µ*L RPMI medium, smashed again using a pestle, centrifuged (1,000 rpm, 3 min, RT) and the supernatant was pooled with the first one. A 1:10 dilution of the sporozoite solution was counted in a Neubauer counting chamber and 10,000 salivary gland sporozoites were used to infect 1 well. After 1.5 hours wells were washed twice with complete DMEM medium and HepG2 cells were allowed to grow in complete DMEM medium supplemented with 1x Antibiotic-Antimycotic (Thermo Fisher Scientific). At 24 h and 48 h post infection cells were fixed using ice cold methanol for 10 min at RT, followed by blocking with 10% FBS/PBS overnight at 4°C. Staining with primary antibody α-*Pb*HSP70 1:300 in 10% FBS/PBS for 2h at 37°C was succeeded by two washing steps with 1% FBS/PBS. Incubation with secondary antibody α-mouse Alexa Fluor 488 1:300 in 10% FBS/PBS was performed for 1h at 37°C. Hoechst 33342 was added and incubated for 5 min at RT followed by two washing steps with 1% FBS/PBS. The assay was mounted in 50% glycerol and sealed using a glass cover slip. Samples were imaged with an Axiovert 200M (Zeiss) microscope and subsequently analysed using ImageJ (Schindelin et al. 2012). In brief, the perimeter of single liver stages was encircled, and the area measured using the internal measurement tool.

### Sporozoite invasion assay

Two days prior to the experiment 180,000 HepG2 cells/well were seeded in an 8-well Permanox Lab-Tek chamber slide (Nunc). Sporozoites were isolated as described above and 10,000 sporozoites/well were used to infect HepG2. At 1.5 h post infection cells were washed twice with complete DMEM medium and fixed using 4% PFA/PBS 20 min at RT. Blocking was performed with 10% FBS/PBS o/n at 4°C followed by incubation with primary antibodies α-*Pb*CSP 1:100 in 10% FBS/PBS (2h 37°C), two washing steps with 1% FBS/PBS and incubation with secondary antibodies α-mouse Alexa Fluor 488 1:300 in 10% FBS/PBS (1h 37°C). After two washing steps with 1% FBS/PBS, cells were permeabilized by addition of ice cold methanol and incubation at RT for 10 min. Blocking with 10% FBS/PBS (4°C over night) was followed by an incubation with primary antibodies α-*Pb*CSP 1:100 in 10% FBS/PBS (2h 37°C), two washing steps with 1% FBS/PBS and incubation with primary antibodies α-mouse Alexa Fluor 546 1:300 in 10% FBS/PBS (1h 37°C). The assay was mounted in 50% glycerol after two washing steps with 1% FBS/PBS.

### Measuring organ-specific parasite load

To measure the parasite load of different organs C57BL/6 mice were infected by intravenous injection of 20,000 salivary gland sporozoites. At 42h after infection organs were harvested, homogenized and the RNA was isolated using Trizol according to the manufacturers protocol. Isolated RNA was treated with DNase using the Turbo DNA-free^TM^ Kit (Invitrogen). Subsequently RNA content of generated samples was measured using a NanoDrop and liver, intestine, spleen and lung samples were pooled in equal amounts of RNA to generate single samples for each parasite line and harvested organ. RNA pools were used to synthetize cDNA using the First Strand cDNA Synthesis Kit (Thermo Fisher Scientific). Quantitative RT-PCR was performed in triplicates on an ABI7500 (Applied Biosystems) using a 2x SYBR green Mastermix (Applied Biosystems). *Plasmodium berghei* 18S rRNA was used to quantify parasites and mouse-specific GAPDH was utilized as housekeeping gene for normalization. Subsequently the ΔcT was plotted as mean of all replicates per parasite strain and harvested organ (Schmittgen & Livak 2008).

### Ethics statement

All animal experiments were performed according to the German Animal Welfare Act (Tierschutzgesetz) and executed following the guidelines of the Society of Laboratory Animal Science (GV-SOLAS) and of the Federation of European Laboratory Animal Science Associations (FELASA). All experiments were approved by the responsible German authorities (Regierungspräsidium Karlsruhe).

### Animal work

For all experiments female 4-6 week old Naval Medical Research Institute (NMRI) mice or C57BL/6 mice obtained from Janvier laboratories were used. Transgenic parasites were generated in the *Plasmodium berghei* ANKA background (Vincke & Bafort 1968) either directly in wild-type or from wild-type derived strains (e.g. *trap(-)rec* and *fluo*). Parasites were cultivated in NMRI mice while transmission experiments with sporozoites were performed in C57Bl/6 mice only.

### Statistical analysis

Statistical analysis was performed using GraphPad Prism 5.0 (GraphPad, San Diego, CA, USA). Data sets were either tested with a one-way ANOVA or a Student’s t test. A value of p<0.05 was considered significant.

## Acknowledgements

We thank Christian Sommerauer for rearing *Anopheles stephensi* mosquitoes as well as Christine Hopp for helpful discussions at the BioMalPar conference. This work was funded by the Human Frontier Science Program (RGY0071/2011), the German Research Foundation (SFB 1129), NIH Grant CA 31798, and the European Research Council (ERC StG 281719). DK was a member of the Hartmut Hoffman-Berling International Graduate School (HBIGS). FF is member of CellNetworks Cluster of excellence at Heidelberg University. We acknowledge the microscopy support from the Infectious Diseases Imaging Platform (IDIP) at the Center for Integrative Infectious Disease Research.

## Supplemental figures

**Figure S1.**
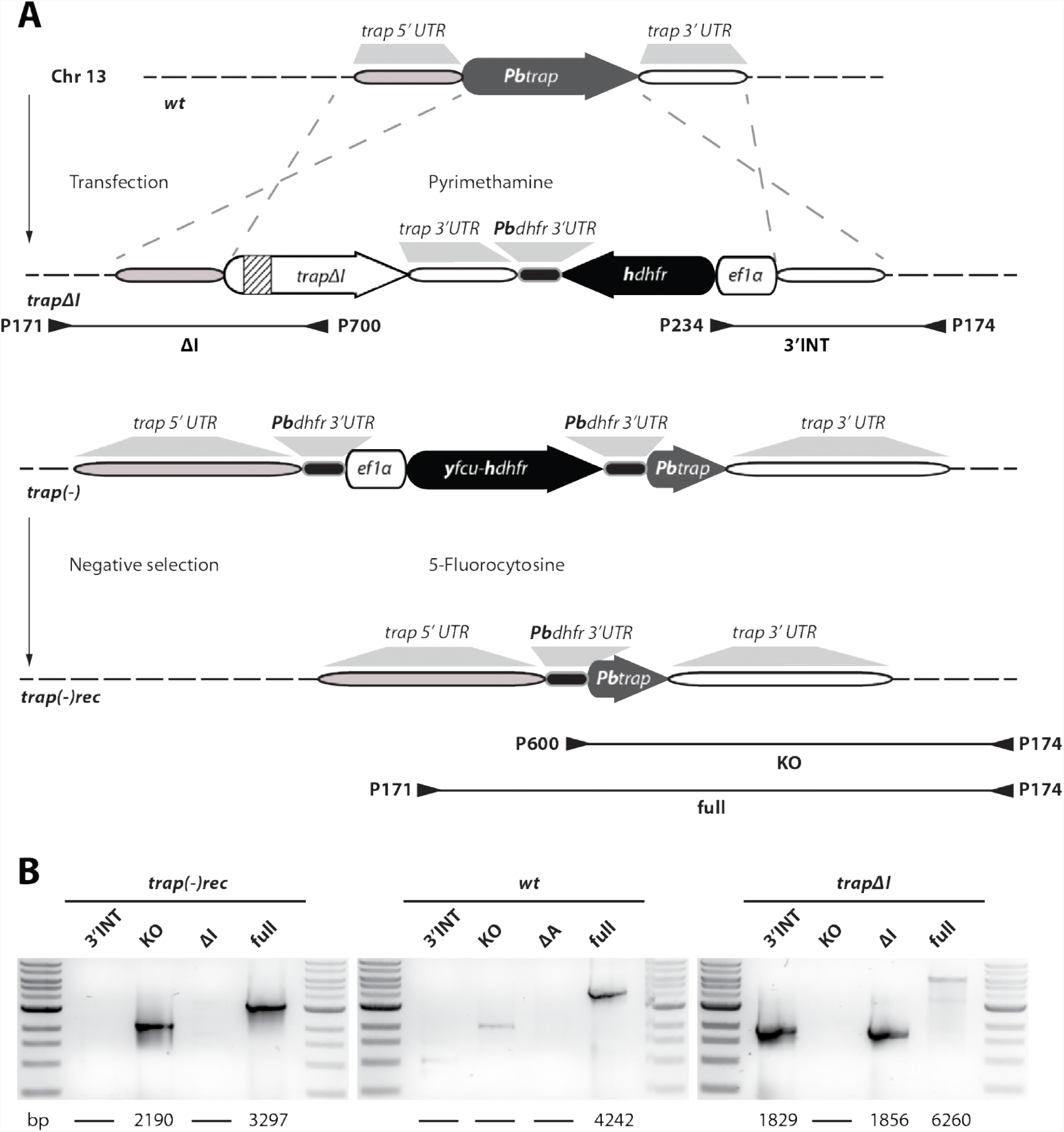
Generation of *trapΔI* and *trap(-)rec* parasites. **A)** Strategy to generate parasites that lack the I-domain of TRAP (*trapΔI*) or a *trap(-)* line without resistance marker (*trap(-)rec)* by double crossover homologous recombination. For the generation of *trapΔI* parasites the *trap* locus in wild-type (*wt*) parasites was replaced with a gene copy that lacks the I-domain including a positive resistance marker (*hdhfr*) driven by the *ef1α* promoter. Note that the inserted copy was codon modified for *E. coli K12* (the codon modified gene copy is indicated in white while the wild-type allele is shown in grey) to avoid unwanted crossover events with the C-terminus of the *trap* open reading frame (ORF). The deleted sequence encoding the I-domain is indicated by stripes. The TRAP knockout was generated with a Plasmogem vector (see material & methods) by replacing the *trap* locus with a selection cassette that enables positive and negative selection. The resulting *trap(-)* parasites were treated with 5-fluorocytosine to select for parasites that lost the resistance cassette (*trap(-)rec*). Primer and predicted length of PCR products are indicated by arrowheads and black lines below the scheme. **B)** PCR analysis of isogenic populations of *trapΔI*, *trap(-)rec* in comparison to wild-type *(wt)*. Sizes of PCR products are indicated below the images.

**Figure S2.**
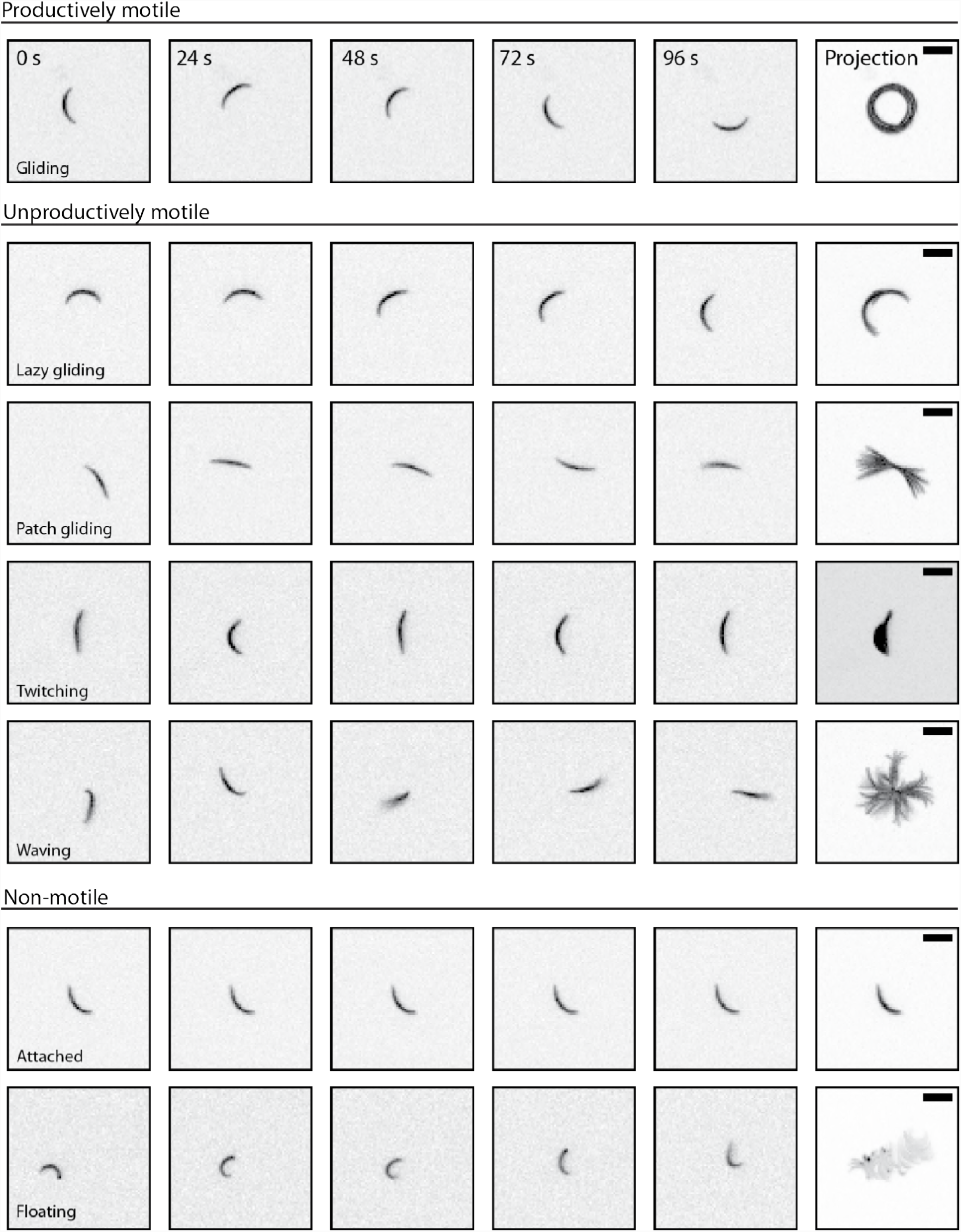
Movement patterns exhibited by sporozoites. Sporozoites placed on solid substrates can exhibit different types of movement as shown in the figure. Gliding, sporozoites glide continuously in circles. Lazy gliding, sporozoites move in a circular manner but never complete a full circle within five minutes. Patch gliding, sporozoites glide back and forth over a single adhesion site. Twitching, sporozoites are attached to the surface and bend back and forth continuously. Waving, sporozoites are attached on one end and turn their body around this adhesion. Attached, sporozoites are completely attached but not moving. Floating, sporozoites are not attached and drift with the flow of the medium. Gliding results in directed forward movement and was classified as productive motility. Other types of movements (lazy gliding, patch gliding, twitching, and waving) lead to persistence of sporozoites at a single site and were classified as unproductive motility. Attached and floating sporozoites were classified as non-motile. Scale bar: 10 *µ*m.

**Figure S3.**
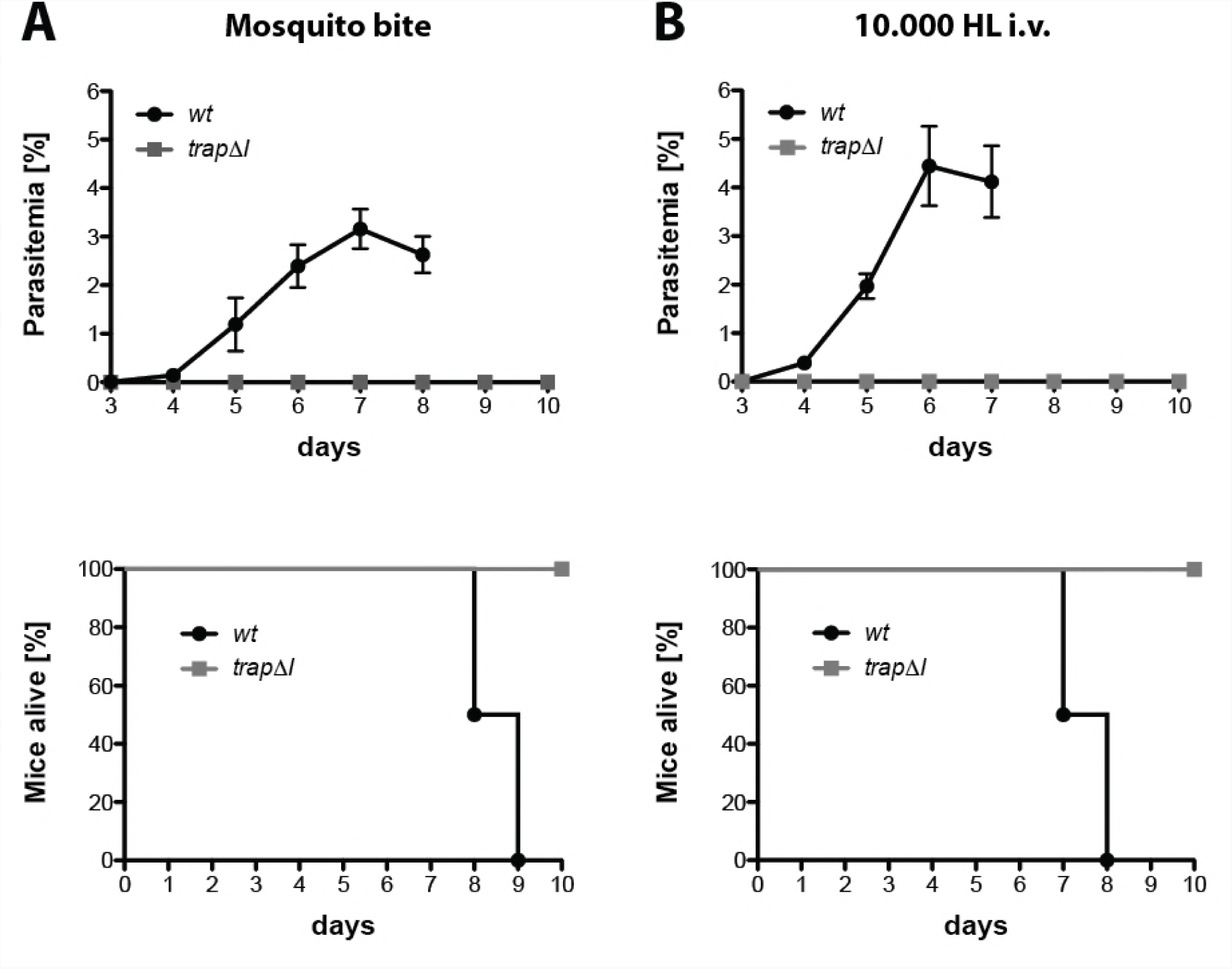
*trapΔI* sporozoites are not infective to mice. Mice were either exposed to 10 infected mosquitoes **(A)** or injected intravenously with 10,000 Hemolymph (HL) sporozoites **(B)** of *trapΔI* or wild-type *(wt)*. Parasite growth and survival of infected mice was monitored for 10 days post infection. Survival graphs correspond to the growth curve shown above. Shown is the mean parasitemia ± SEM of four mice for each parasite line.

**Figure S4.**
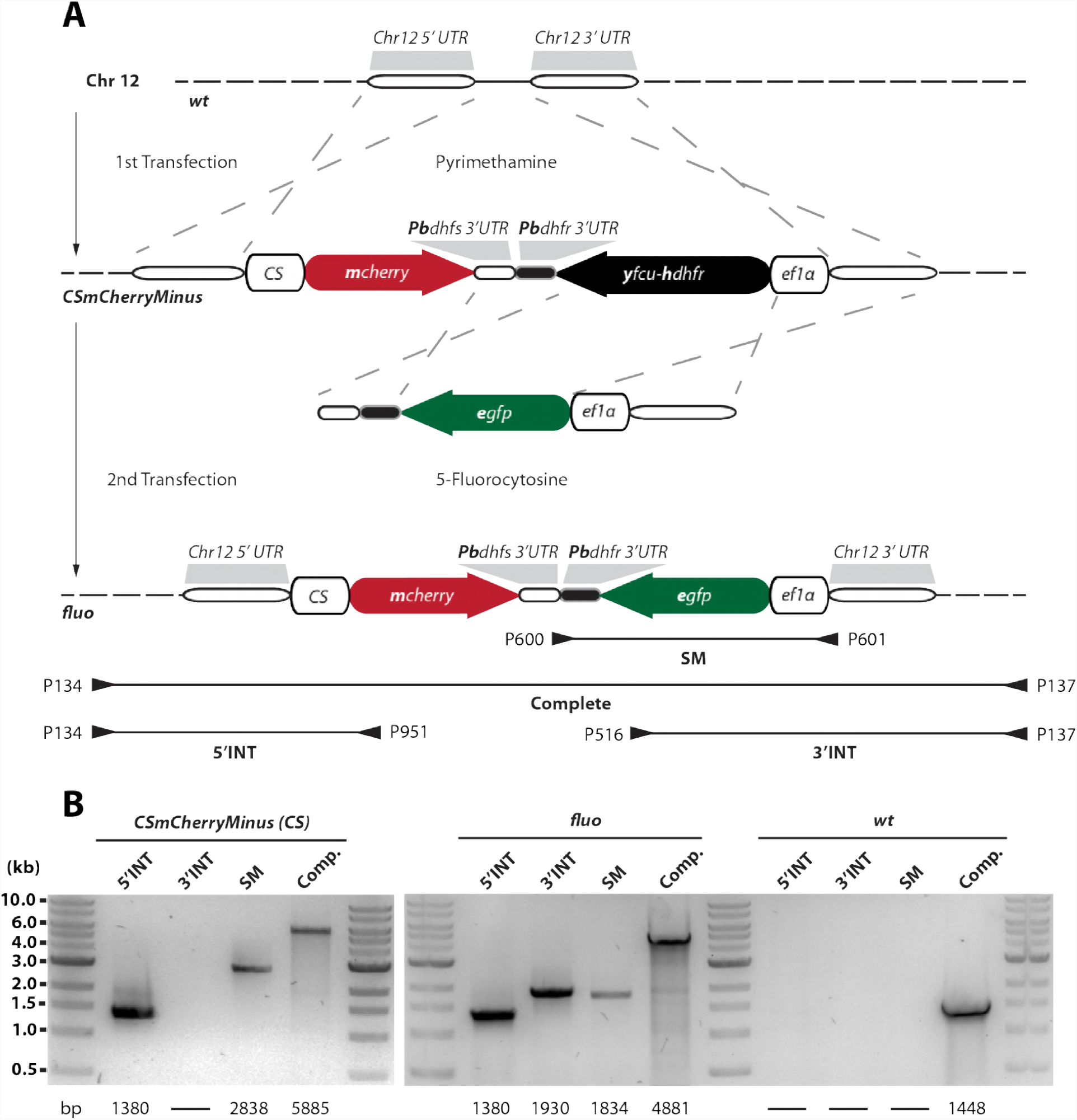
Generation of the selection marker free reporter line *fluo*. **A)** Illustration showing the integration via double crossover homologous recombination of the mCherry reporter cassette (Pb262) and the positive-negative selection marker *yfcu-hdhfr* into chromosome 12 to generate the recipient line *CSmCherryMinus*. This line does not contain a second *3’dhfr* sequence downstream of the *ef1α* promoter that is needed to recycle the selection cassette by negative selection. Instead this line can be used as recipient line to integrate DNA sequences that can replace the selection cassette. The presence of the *ef1α* promoter is important for a downstream gene (Klug et al. 2016). The *fluo* reporter line was created in a second transfection by integration of a DNA sequence containing a short sequence of chromosome 12 as well as the *egfp* gene driven by the *ef1α* promoter and the *3’dhfr* terminator. This sequence replaced the selection cassette by double crossover homologous recombination and gave rise to the *fluo* reporter line. The line was selected with 5-fluorocytosine and cloned as described in material & methods. Location and number of primers used for genotyping are indicated below the scheme. Note that the illustration is not drawn to scale. **B)** Genotyping via PCR shows positive integration of the *mCherry* gene (5’INT) and the *yfcu-hdhfr* selection marker (SM) into chromosome 12 for *CSmCherryMinus*. The PCR for the 3’integration (3’INT) targets the *egfp* gene, which is absent in *CSmCherryMinus* and therefore negative. PCR analysis of the *fluo* line shows presence of *mcherry* (5’INT) and *egfp* (3’INT). The PCR for the *yfcu-hdhfr* selection marker is still positive because the used primers bind to flanking regions, which are still present in the *fluo* line but loss of the *yfcu-hdhfr* gene is indicated by a shift in size. The wild-type (*wt*) control shows no product for PCRs targeting sequences within the integrated sequence. Compared to *CSmCherryMinus* and *fluo* parasites the PCR that amplifies the complete locus (Comp.) results in a much smaller product for wild-type (*wt*) indicating no integration in this site.

**Figure S5.**
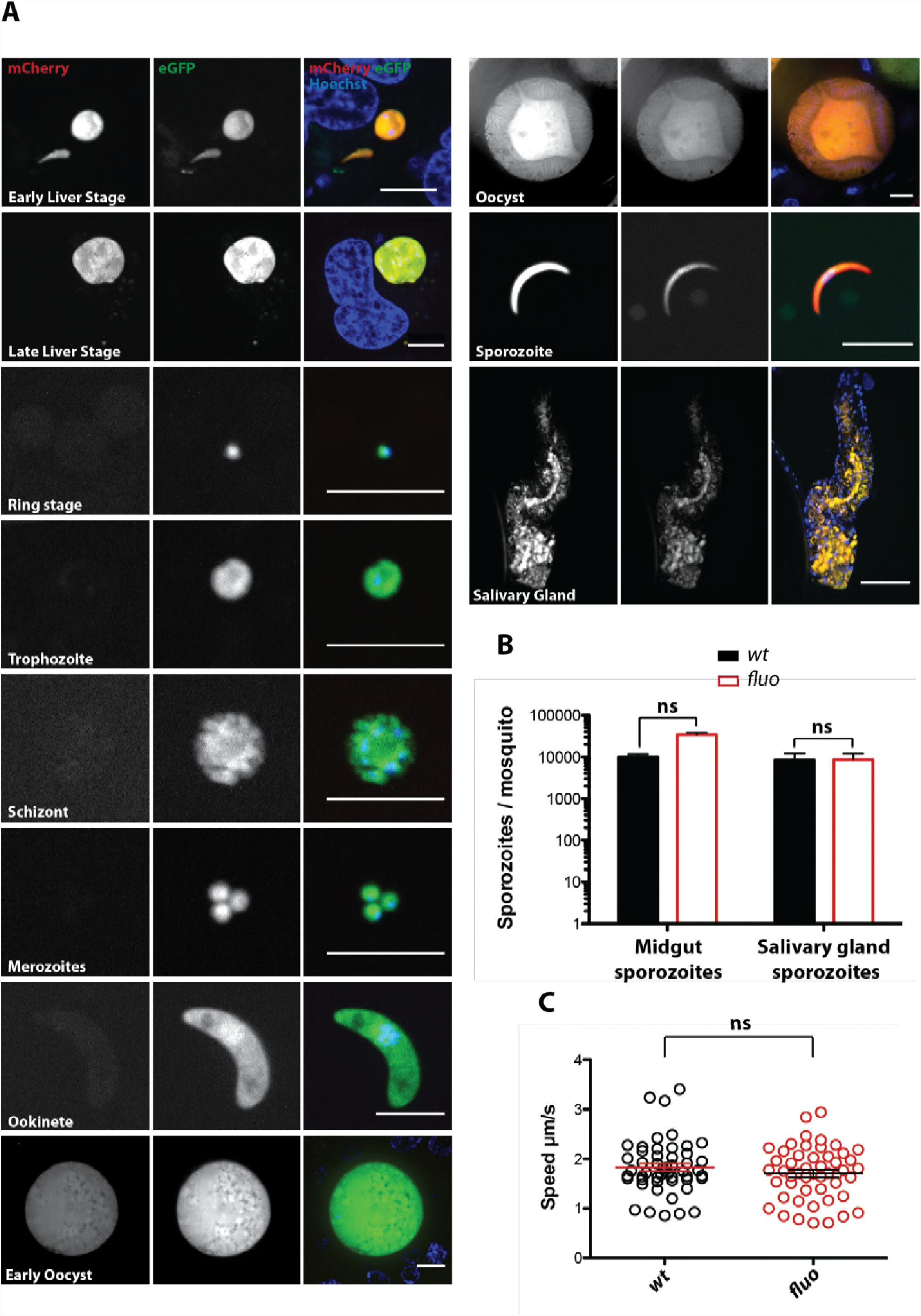
Phenotypic characterization of the *fluo* line. **A)** Live imaging of the *fluo* line across the life cycle revealed constitutive expression of eGFP in all evaluated stages as well as strong mCherry expression in late oocysts, sporozoites and liver stages. Shown are all stages except zygotes as well as female and male gametocytes. Parasites were additionally stained with Hoechst 33342 to visualize DNA. Scale bar for all images except salivary glands: 10 *µ*m. Scale bar for image with salivary gland: 100 *µ*m. Shown are single images or maximum projections of stacked images (salivary gland). **B)** Sporozoite numbers of midgut and salivary gland sporozoites in mosquitoes infected with *fluo* and wild-type (*wt*) parasites. Shown are ≥ 3 counts from three different feeding experiments. Data were tested for significance using Mann-Whitney test. **C)** The speed of *fluo* sporozoites is comparable to wild-type (*wt*). Shown are the average speeds of 50 salivary gland sporozoites per strain that were tracked for 2 minutes and 30 seconds. Horizontal lines indicate the overall mean ± SEM. Data were tested for significance with the Mann-Whitney test.

**Figure S6.**
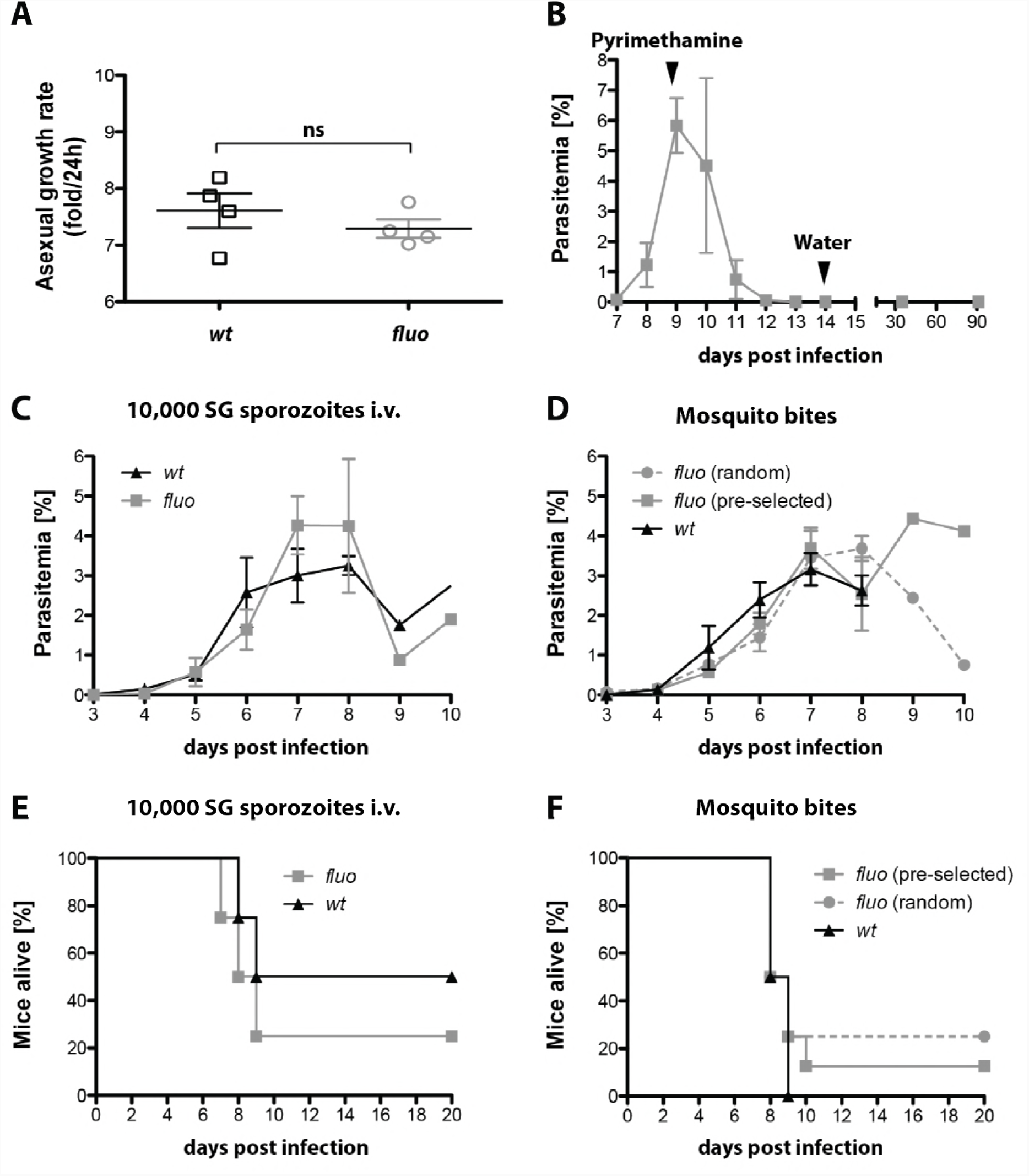
The *fluo* line is sensitive to pyrimethamine and shows similar transmission efficacy and blood stage growth as wild-type (*wt*). **A)** The *fluo* line shows no difference in blood stage growth compared to wild-type (*wt*). Naive mice were infected by intravenous injection of 100 blood stage parasites and parasitemia was monitored daily by Giemsa stained blood smears. Blood stage growth was calculated based on parasitemia on day 9 post infection (Klug et al. 2016). Shown is the mean ± SEM of four mice per parasite line. Data were tested for significance with the Mann-Whitney test. **B)** The *fluo* line is sensitive to pyrimethamine. Two mice were infected with single blood stage parasites. On day 9 post infection pyrimethamine was given within the drinking water. The development of parasitemia was monitored for 91 days, administration of pyrimethamine and drug-free drinking water are indicated with arrowheads within the graph. Shown is the mean parasitemia ± SEM. **C)** Mice were infected with the *fluo* line or wild-type (*wt*) by intravenous injection of 10,000 salivary gland sporozoites or by bite of infected mosquitoes. Shown is the mean parasitemia ± SEM of four mice per group. **(D)**. Infected mosquitoes used for *in vivo* experiments were either pre-sorted for fluorescent parasites in the midgut (*fluo* pre-selected) or randomly selected (*fluo* random and *wt*). **E)** and **F)** Survival of mice infected intravenously with 10,000 salivary gland sporozoites or by mosquito bites corresponding to the growth curves shown above in panels C and D, respectively. Survival of infected mice was monitored for 20 days post infection.

**Figure S7.**
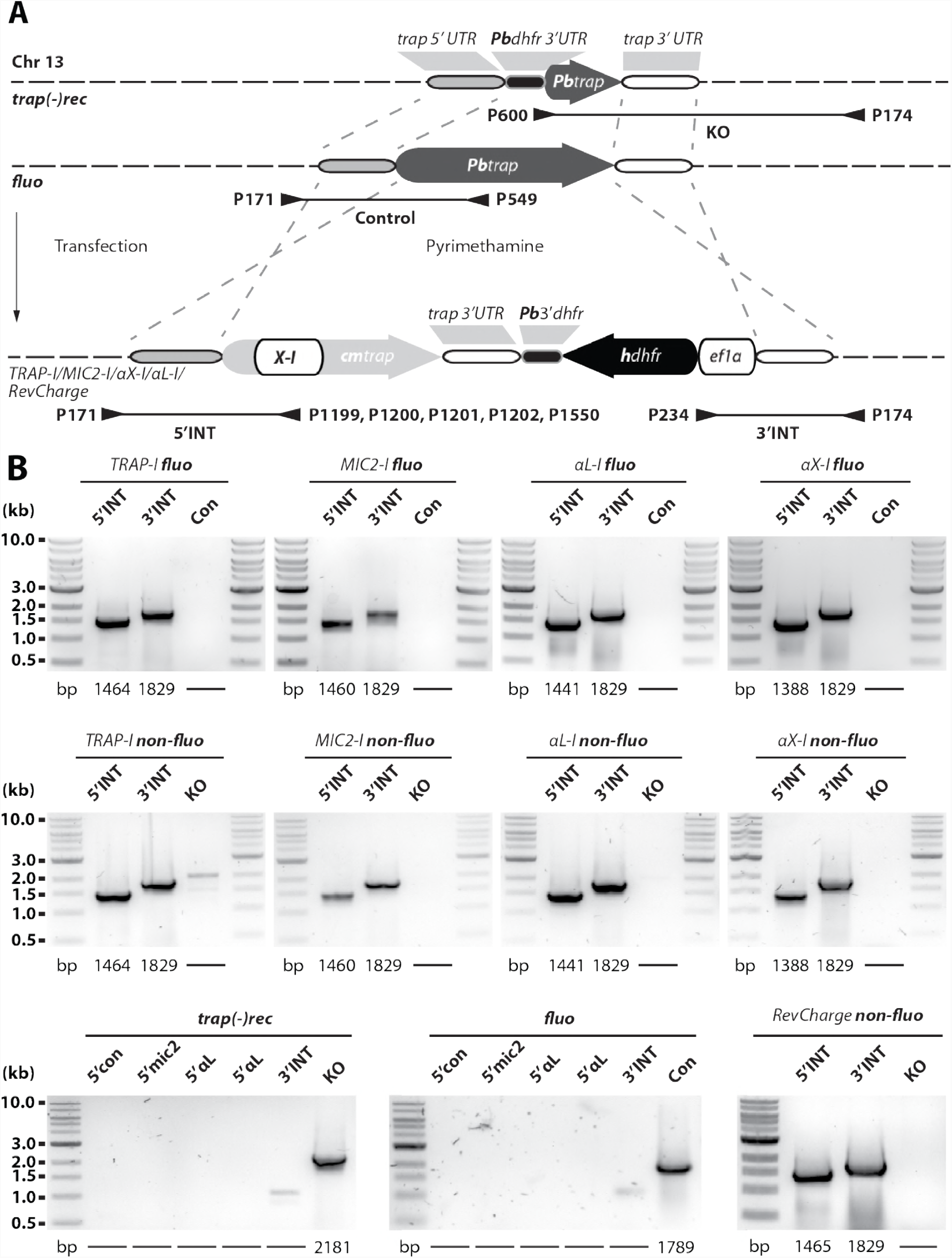
Generation of *P. berghei* strains expressing TRAP with different I-domains. **A)** *trap* genes with exchanged I-domains were transfected into *fluo* and *trap(-)rec* parasites to generate two sets of mutants. Transgenic parasites were generated by double crossover homologous recombination replacing the endogenous *trap* gene (*fluo line*) or complementing the *trap* locus in *trap(-)rec* parasites. Five different parasite lines were generated: *TRAP-I*, expressing the wild-type allele of TRAP and thus serving as a control line; *MIC2-I*, chimera expressing the I-domain of micronemal protein 2 (MIC2) from *Toxoplasma gondii*; *αX-I*, chimera expressing the I-domain of the human integrin αX; *αL-I*, chimera expressing the I-domain of the human integrin αL; *RevCharge*, wild-type allele containing seven mutations that shift the surface charge of the I-domain from a pI of 9.7 to 6.8 (see **Figure 6A**). The *trap* coding sequence in all generated lines was codon modified for *E. coli K12* to avoid unwanted crossover events and to distinguish chimeras from wild-type (*wt*). Binding sites and numbers of primers used for genotyping as well as the length of amplified PCR products are indicated with arrowheads and black lines below the scheme. Note that the illustration is not drawn to scale. **B)** To control for correct integration of the transfected DNA sequences, three different PCRs were performed. The 5’INT PCR amplifies the 5’ end of the integrated sequence with a primer that binds upstream of the integration site matching a primer that binds specifically to the sequence encoding the I-domain. The 3’INT PCR amplifies the 3’ end of the integrated sequence with a primer that binds downstream of the integration site matching a primer in the selection cassette. The control PCR (Con or KO) uses primers that are specific for the recipient lines *fluo* or *trap(-)rec*. The length of the expected PCR products is depicted below the gel images. Shown are only PCR results of isogenic populations cloned by limiting dilution.

**Figure S8.**
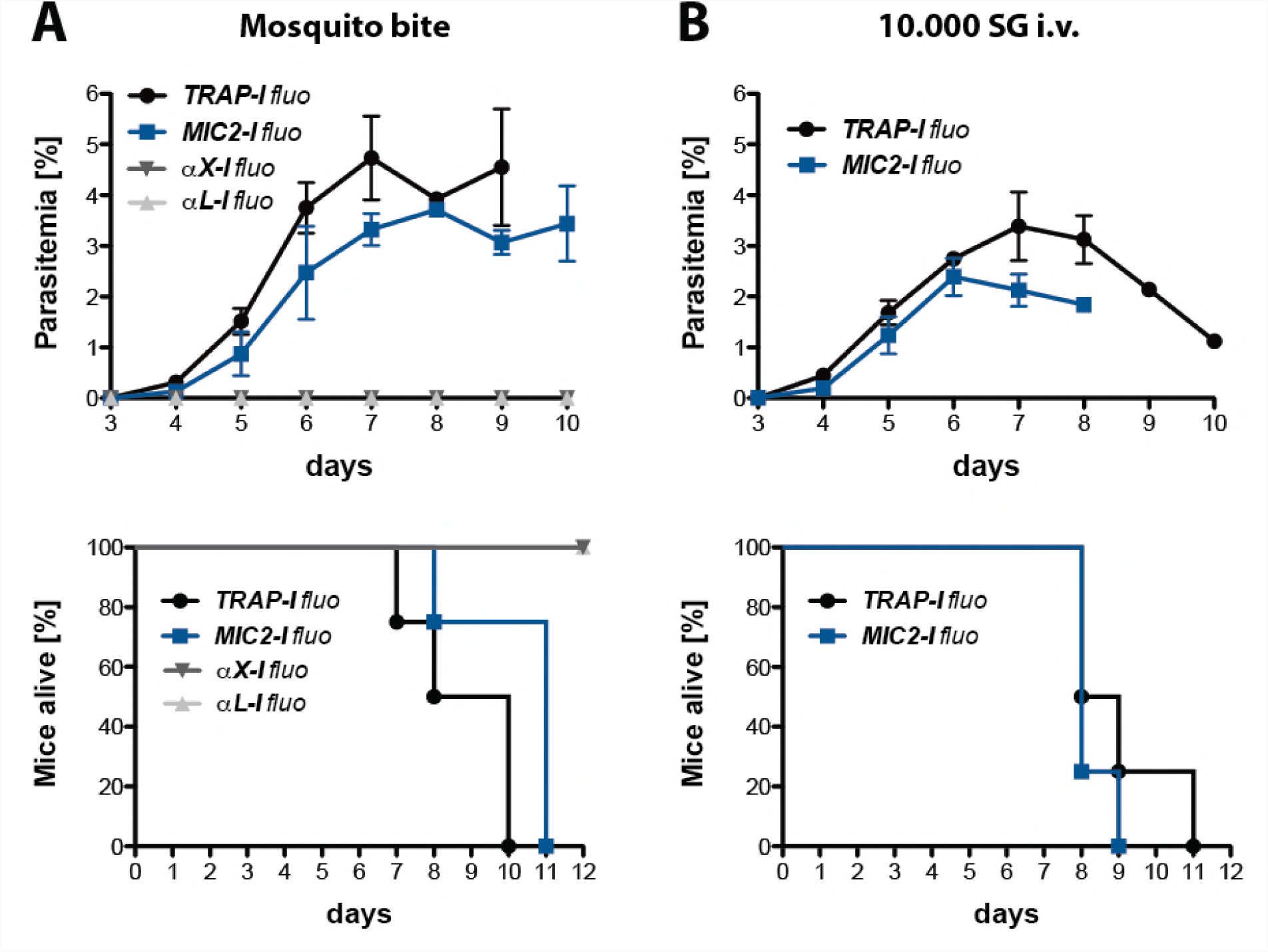
Transmission efficacy of SG sporozoites expressing TRAP with different I-domains transmitted either by mosquito bite or by intravenous injection. Mice were either exposed to 10 infected mosquitoes **A)** or injected intravenously with 10,000 salivary gland (SG) sporozoites **B).** Shown is the parasitemia of four infected mice per parasite line as mean ± SEM from day 3 to day 10. The survival of infected mice was monitored for 12 days post infection. The survival graphs correspond to the growth curves shown above. Per parasite line and experiment four mice were infected. Note that only growth curves and survival graphs of fluorescent parasite lines are shown.

**Figure S9.**
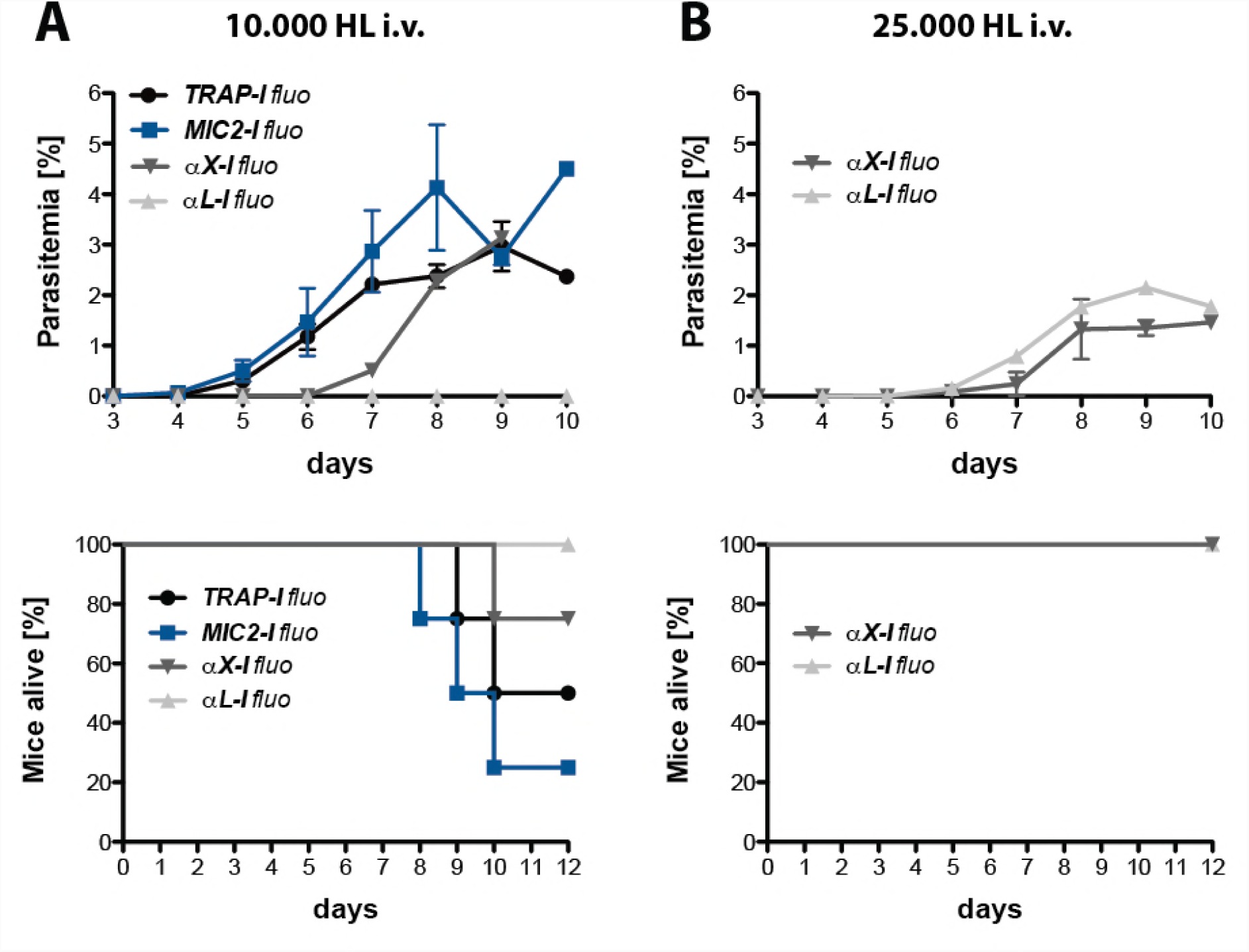
Transmission efficacy of intravenously injected HL sporozoites expressing TRAP with different I-domains. Mice were either injected intravenously (i.v.) with 10,000 **A)** or 25,000 **B)** hemolymph (HL) sporozoites. Injections with 25,000 HL sporozoites were only performed with *αX-I fluo* and *αL-I fluo* parasites. Shown is the parasitemia of four infected mice per parasite line as mean ± SEM from day 3 to day 10. The survival of infected mice was monitored for 12 days post infection. Survival graphs correspond to the growth curves shown above. Per parasite line and experiment four mice were infected. Note that only growth curves and survival graphs for fluorescent parasite lines are shown.

**Figure S10.**
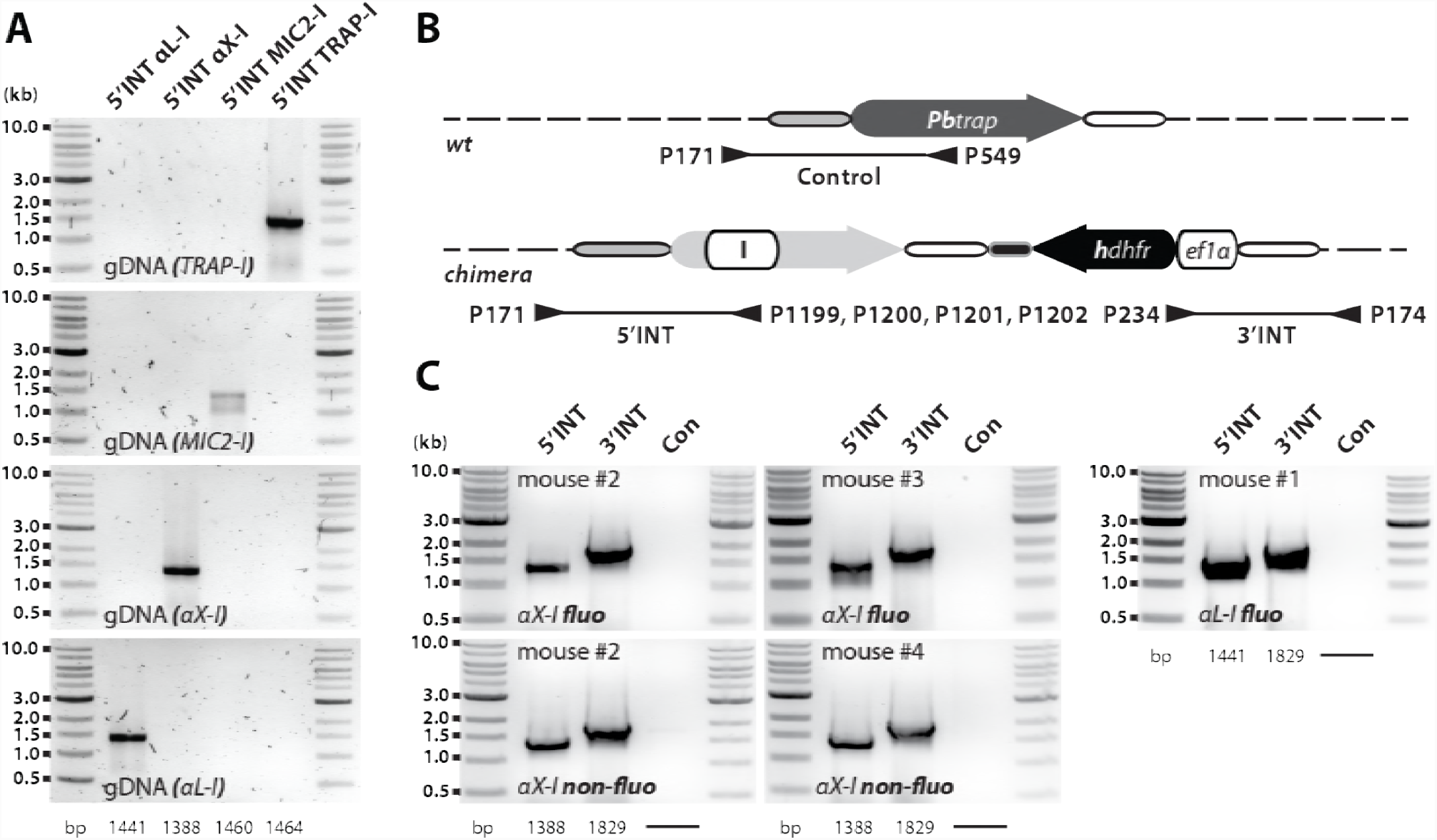
Sporozoites expressing the integrin I-domains αX or αL are infective to mice. **A)** Primers were designed to bind specific to sequences encoding for the different I-domains TRAP-I, MIC2-I, αX-I and αL-I. **B)** The genotype of parasites isolated from infected mice was determined by PCR with three different primer combinations: control (con); amplification of the 5’UTR including the N-terminal end of the TRAP wild-type ORF, 5’INT; amplification of the 5’UTR including the N-terminal end of the codon modified TRAP ORF, 3’INT; amplification of the 3’UTR including the C-terminal part of the selection cassette. Illustration is not drawn to scale. **C)** Mice who became blood stage positive after injection of 10,000 or 25,000 α*X-I* or α*L-I* HL sporozoites were genotyped via PCR. In addition, sequencing of the *trap* locus revealed the correct identity of the used parasite line in all tested mice (data not shown). Note that two mice infected with α*X-I* sporozoites died due to cerebral symptoms and could not be genotyped.

**Figure S11.**
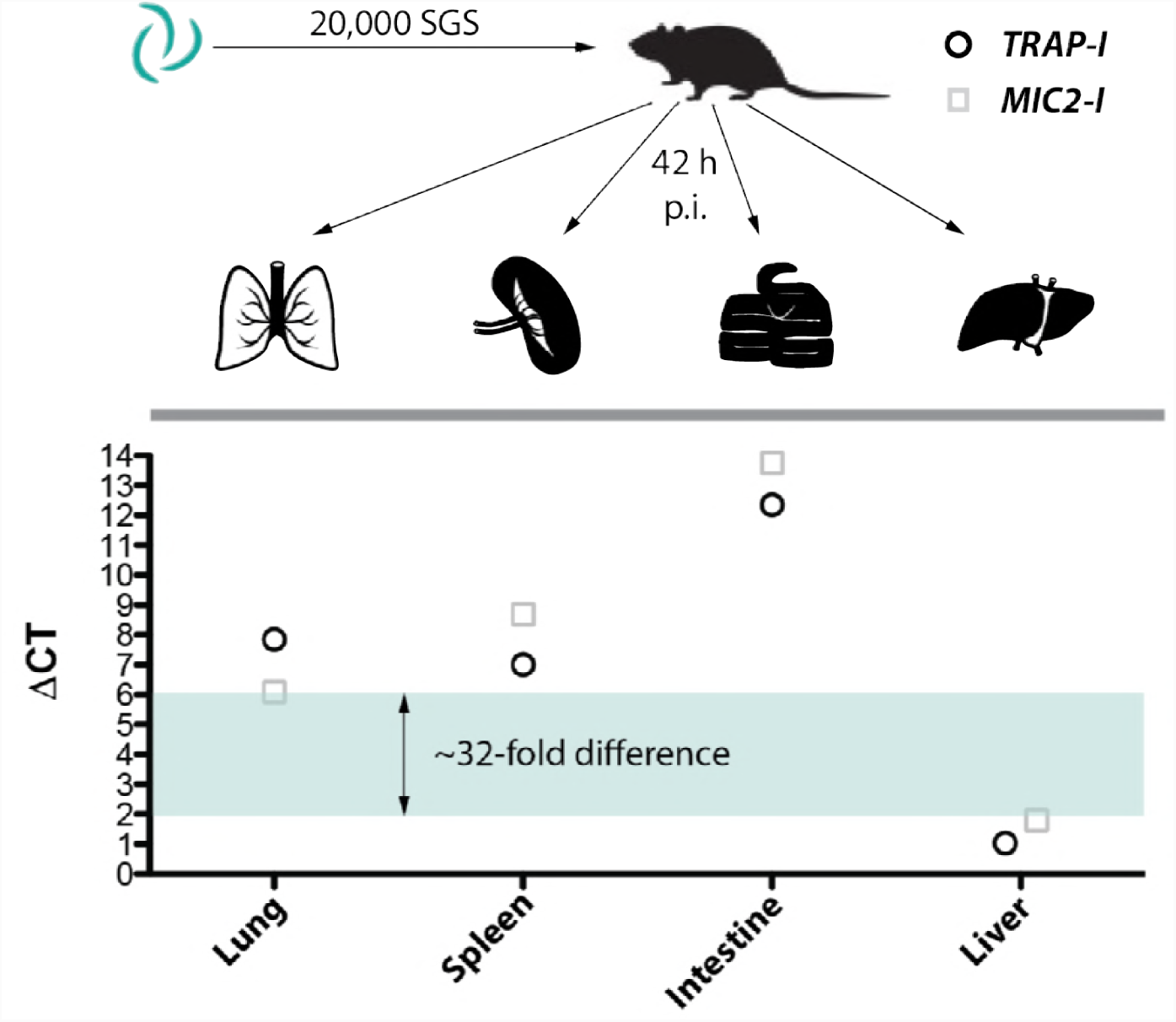
*MIC2-I* sporozoites show no altered tissue tropism compared to *TRAP-I*. Parasite load in lung, spleen, small intestine and liver of mice infected with 20,000 *MIC2-I* or *TRAP-I* salivary gland sporozoites (SGS). Organs were harvested 42 h post infection to isolate RNA. Per organ and parasite line equal amounts of RNA were pooled from four infected mice to generate one *TRAP-I* and one *MIC2-I* sample per harvested organ. qRT-PCR was performed in technical triplicates and the mean was plotted as single ΔCT value. For further details please see material & methods.

**Data S1.**
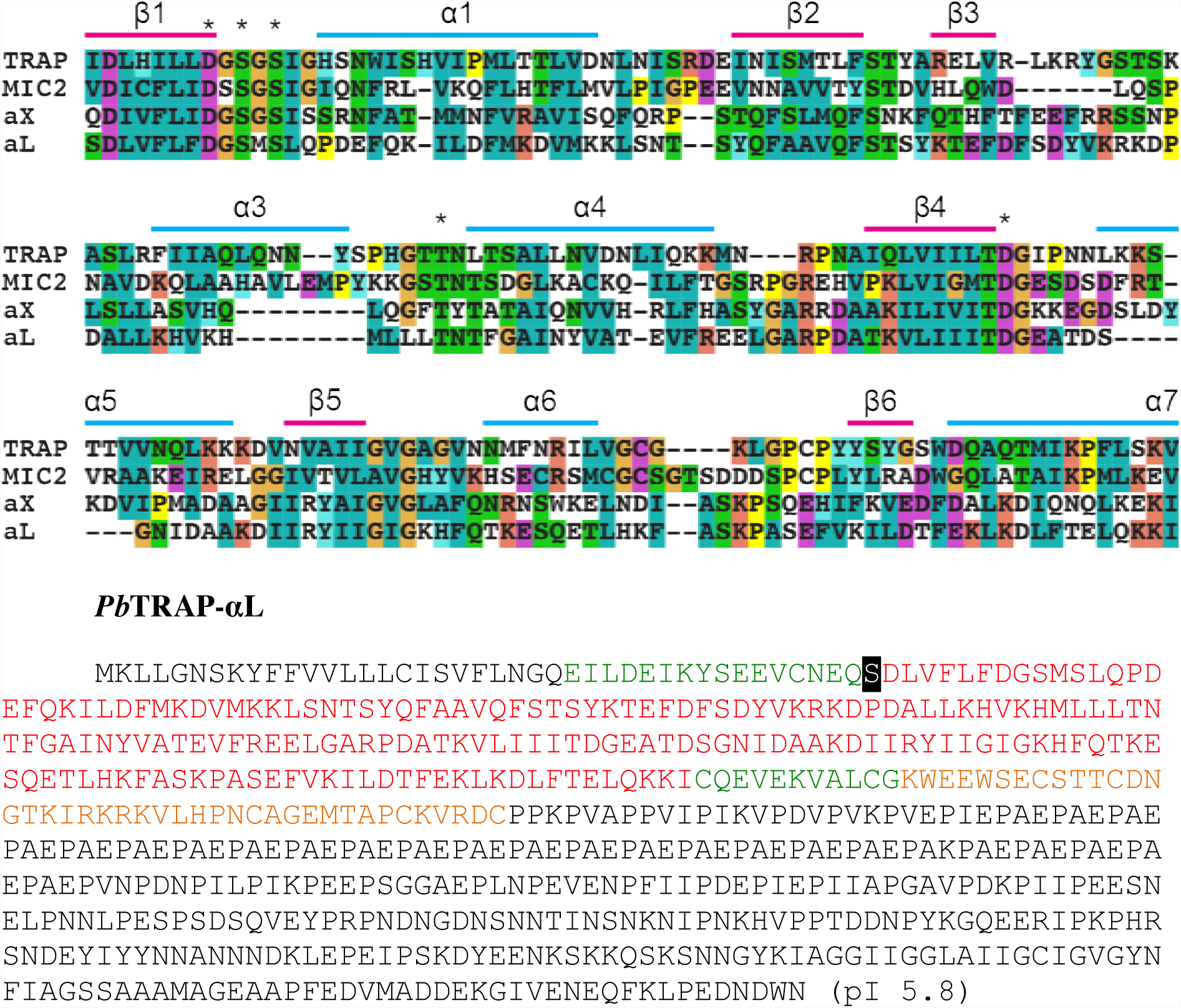
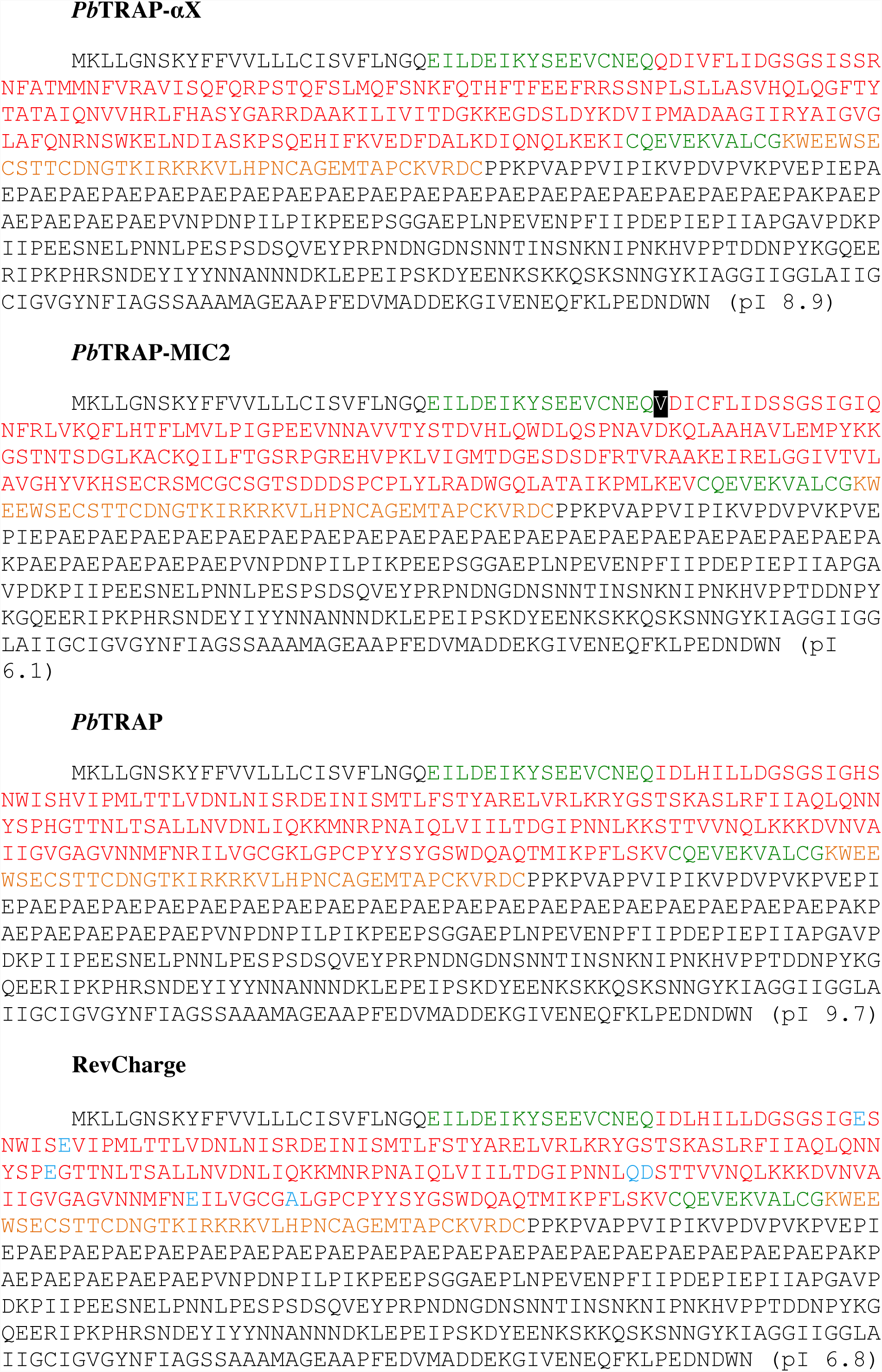
Alignment of the I-domain portions exchanged in this study and the amino acid sequences of the TRAP variants expressed by the parasite lines *TRAP-I*, *MIC2-I*, *αX-I, αL-I, and RevCharge*. The portions of the four I-domains that were exchanged were aligned by sequence and structure (Song et al. 2012; Song & Springer 2014). Secondary structure elements are named and MIDAS residues are asterisked above the alignment. Below the alignment are shown the sequences of each TRAP replacement. Residues that are part of the extendable ß-ribbon are written in green, residues that form the remainder of the I-domain are written in red, residues of the thrombospondin domain are written in orange, and the remaining native residues of *Pb*TRAP are written in black. Residues written in blue were introduced into wild-type *Pb*TRAP to generate a more negative charge on the portion of the I-domain surface surrounding the MIDAS in the *RevCharge* mutant. Residues written in white on a black background were mutated to create a better fitting of the exchanged portion of the I-domain with the N-and C-terminal segments of the *Pb*TRAP I-domain/extendable ß-ribbon. The calculated pI of the I-domain region is shown in parentheses.

**Table S1.**
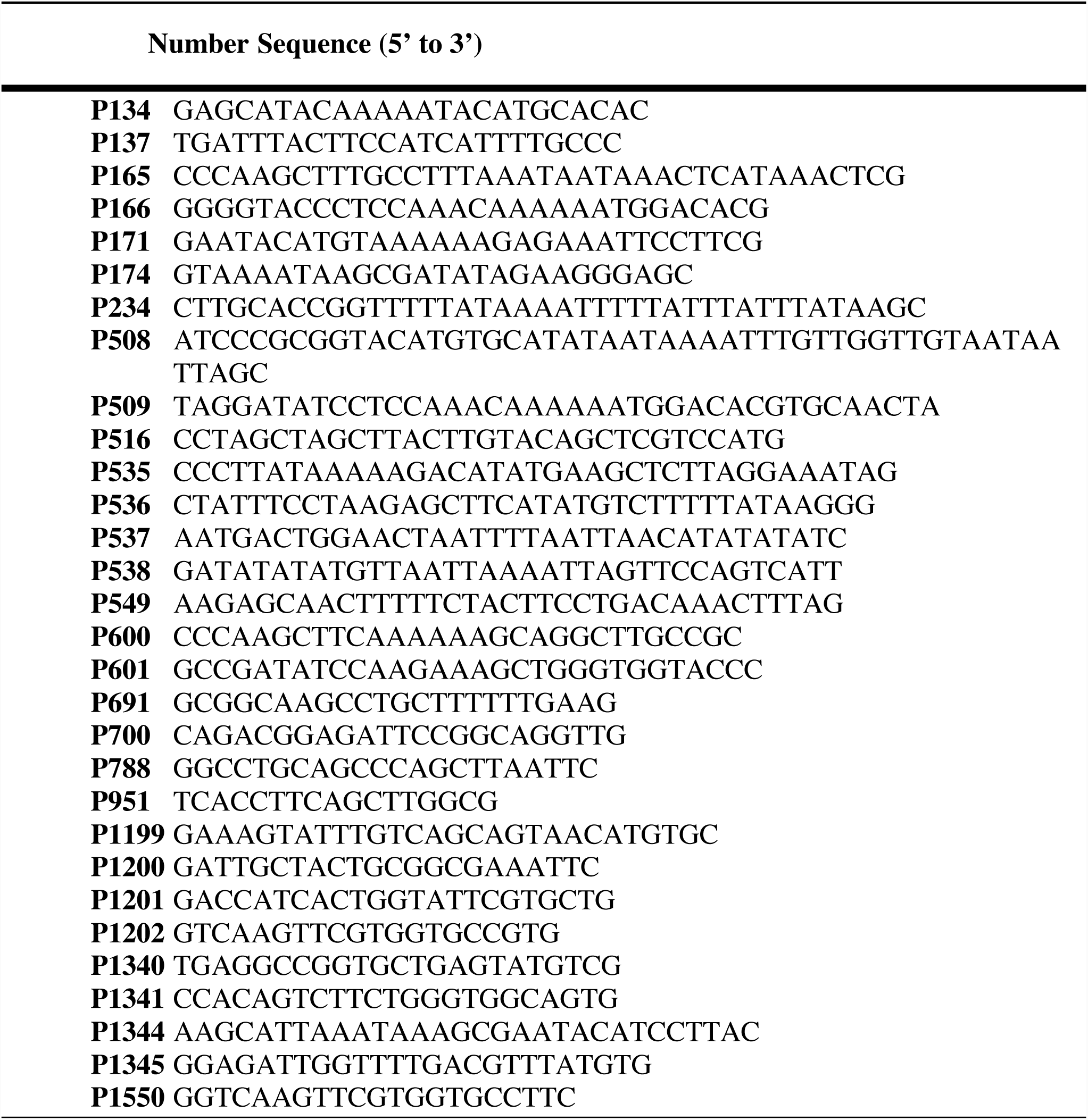
Primer sequences. Primers used for the generation and genotyping of the parasite lines presented in this study.

**Movie S1. Z-projection through salivary gland infected with *aX-I fluo* sporozoites.**

Shown is an image series in Z-direction of a salivary gland infected with *aX-I fluo* sporozoites. Images were taken on an Axiovert 200M (Zeiss) with a 63x (N.A. 1.3) objective. study.

